# A likelihood approach for uncovering selective sweep signatures from haplotype data

**DOI:** 10.1101/678722

**Authors:** Alexandre M. Harris, Michael DeGiorgio

## Abstract

Selective sweeps are frequent and varied signatures in the genomes of natural populations, and detecting them is consequently important in understanding mechanisms of adaptation by natural selection. Following a selective sweep, haplotypic diversity surrounding the site under selection decreases, and this deviation from the background pattern of variation can be applied to identify sweeps. Multiple methods exist to locate selective sweeps in the genome from haplotype data, but none leverage the power of a model-based approach to make their inference. Here, we propose a likelihood ratio test statistic *T* to probe whole genome polymorphism datasets for selective sweep signatures. Our framework uses a simple but powerful model of haplotype frequency spectrum distortion to find sweeps and additionally make an inference on the number of presently sweeping haplotypes in a population. We found that the *T* statistic is suitable for detecting both hard and soft sweeps across a variety of demographic models, selection strengths, and ages of the beneficial allele. Accordingly, we applied the *T* statistic to variant calls from European and sub-Saharan African human populations, yielding primarily literature-supported candidates, including *LCT, RSPH3*, and *ZNF211* in CEU, *SYT1, RGS18*, and *NNT* in YRI, and *HLA* genes in both populations. We also searched for sweep signatures in *Drosophila melanogaster*, finding expected candidates at Ace, *Uhg1*, and *Pimet*. Finally, we provide open-source software to compute the *T* statistic and the inferred number of presently sweeping haplotypes from whole-genome data.

## Introduction

A selective sweep is a genomic signature resulting from positive selection in which the linked variants surrounding the site under selection rise to high frequency together in a population, thereby yielding a footprint of reduced diversity that can span across megabases [Przeworski, 2002, Gillespie, 2004, Kim and Nielsen, 2004, Garud et al., 2015, Hermisson and Pennings, 2017]. Thus, a recent selective event is identifiable in polymorphism data from a region of extended haplotype homozygosity, and the signal of a selective sweep accordingly decays over time as mutation and recombination break up long haplotypes [Sabeti et al., 2002, Schweinsberg and Durrett, 2005, Voight et al., 2006]. Selective sweeps can arise from multiple processes, including the *de novo* emergence of a selectively advantageous allele, selection on standing population haplotypic variation, and recurrent mutation to a selectively advantageous allele [Hermisson and Pennings, 2005, Pennings and Hermisson, 2006a,b]. The former scenario is a hard sweep, in which a single haplotype rises to high population frequency, gradually replacing all other haplotypes as the sweep proceeds to fixation. The latter two scenarios are soft sweeps, in which multiple haplotypes simultaneously rise to high population frequency, and a greater haplotypic diversity underlies the sweep.

Identifying selective sweeps is important because sweeps serve as indicators of recent rapid adaptation in a population, providing insight into the pressures that shaped its present-day levels of genetic diversity [Vatsiou et al., 2016, Librado et al., 2017]. These pressures can vary considerably in their intensity and duration, resulting in selection signals of varying magnitude ranging from prominent, such as *LCT* in Europeans [Bersaglieri et al., 2004], to the more subtle *ASPM*, implicated in the development of human brain size [Kouprina et al., 2004, Peter et al., 2012]. Whereas strong sweeps are typically easy to detect, weaker sweeps typically require a large sample size for detection [Jensen et al., 2007, Pavlidis et al., 2013], and may only be identifiable through sophisticated approaches [Chen et al., 2010]. Selective sweeps, while not the only signature of adaptation in natural populations, are likely to occur at loci where mutations have a large effect size, little negative pleiotropic effects, and contribute to phenotypes that are either monogenic or controlled by few genes [Pritchard and DiRienzo, 2010]. In addition, identifying selective sweeps is important to make inferences about the relative contributions of hard and soft sweeps to adaptive events in study organisms [Garud et al., 2015, Schrider and Kern, 2016, Harris et al., 2018a], which is a topic of continued debate [Jensen, 2014, Schrider and Kern, 2017, Harris et al., 2018b, Mughal and DeGiorgio, 2019].

Multiple powerful methods have been proposed to characterize selective sweeps, and well-established among these are composite likelihood ratio (CLR) methods [Kim and Stephan, 2002, Kim and Nielsen, 2004, Nielsen et al., 2005, Chen et al., 2010, Pavlidis et al., 2013, Vy and Kim, 2015, Racimo, 2016, Huber et al., 2016, DeGiorgio et al., 2016], and haplotype homozygosity-based methods [Voight et al., 2006, Ferrer-Admetlla et al., 2014, Garud et al., 2015, Harris et al., 2018a]. The former category of methods represents approaches in which the probability of neutrality in a genomic region under analysis is compared to the probability of a selective sweep in that region, based on a model of distortion in the site frequency spectrum expected under a sweep. A CLR statistic quantifies support for the alternative hypothesis of selection, with larger values indicating greater support. Although CLR methods make simplifying assumptions in their models [Beaumont et al., 2010, Pavlidis and Alachiotis, 2017], they have demonstrated a powerful capacity for identifying multiple different signatures of selection without the need for computationally intense calculations of full likelihood functions [Kim and Stephan, 2002, DeGiorgio et al., 2014, Huber et al., 2016]. However, because they are typically allele frequency-based approaches, the CLR methods may lack in power to detect soft sweeps in comparison to haplotype-based methods, which can generally detect both [Pennings and Hermisson, 2006b, Ferrer-Admetlla et al., 2014]. Accordingly, the need exists for methods that leverage the power and efficiency of CLR approaches, while providing the sensitivity of haplotype-based approaches.

We introduce an approach for identifying selective sweep signatures using a likelihood ratio framework *T* that is the first haplotype-based method of its kind, intended to address the limitations of previous methods. Our *T* statistic (see *Theory*) has high power to detect recent sweeps from genome-wide polymorphism data and additionally infers the number of presently sweeping haplotypes as a model parameter, providing an additional layer of insight not shared with other CLR methods. This attribute is especially important because it eliminates the need for time- and computation-heavy alternatives, such as training a machine-learning classifier [Lin et al., 2011, Kern and Schrider, 2018, Mughal and DeGiorgio, 2019], or drawing inferences from a posterior distribution by approximate Bayesian computation [Garud et al., 2015, Harris et al., 2018a, Harris and DeGiorgio, 2019]. We demonstrate with simulated data that the *T*-statistic identifies recent hard and soft sweeps, and performs especially well for population size expansion models. As such, our application of the *T* statistic to human and *Drosophila melanogaster* datasets recovered multiple previously-characterized candidate sweeps in both organisms, allowing us to corroborate and enhance our understanding of adaptation in each of their histories.

## Theory

The goal of our approach is to identify genomic signatures of selective sweeps. We achieve this by assigning a *T* statistic to each SNP-delimited window of analysis in the genome. The *T* statistic is a measure of the likelihood that an analysis window is consistent with a selective sweep rather than neutrality. We base this inference on the sample haplotype frequency spectrum, reasoning that a spectrum with few high-frequency haplotypes indicates a sweep, and a spectrum with no moderate- or high-frequency haplotypes indicates neutrality. Thus, our approach is a likelihood ratio test in which the model of neutrality, based on the genome-wide haplotype frequency spectrum, is nested within the model of selection, based on a distortion of the genome-wide haplotype frequency spectrum toward few moderate- or high-frequency haplotypes. We illustrate examples of haplotype frequency spectra for neutrality and sweeps in Figure 1, and also provide a schematic on how key model parameters relate to distortions in the haplotype frequency spectrum.

**Figure 1:**
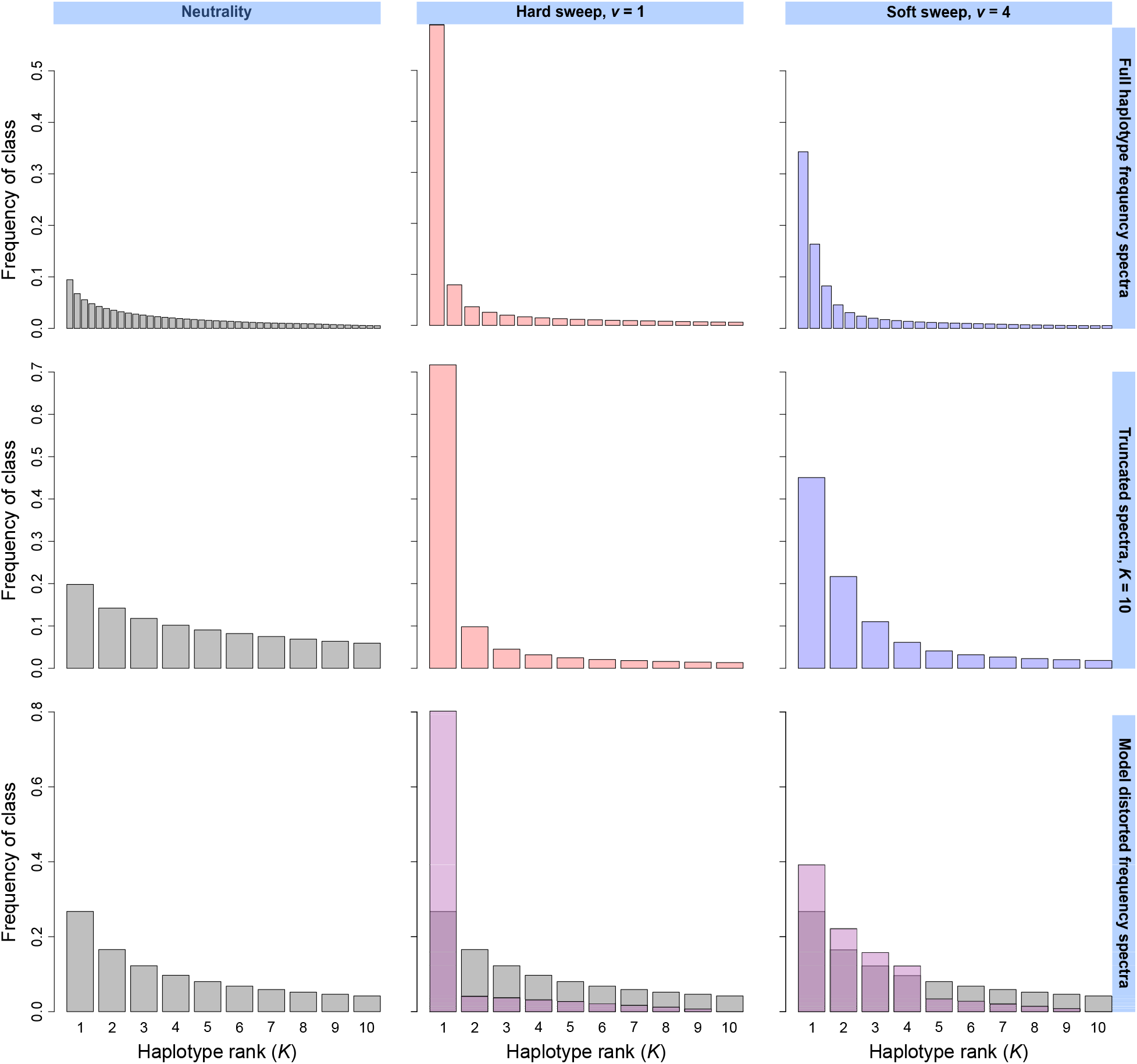
Example simulated haplotype frequency spectra for neutrality, hard sweeps (*ν* =1), and soft sweeps (*ν* = 4). Under neutrality, all sampled haplotypes in an analysis window exist at low frequency, and there are many haplotypes. In contrast, selective sweeps yield high-frequency haplotypes, and fewer total haplotypes (top). Truncated spectra (*K* = 10) preserve their overall shape relative to untruncated spectra above (middle). We distort the truncated neutral spectrum computed from sampled haplotypes to yield spectra corresponding to alternative models (purple), in which the mass of non-sweeping classes is transferred to sweeping classes, resembling the expected pattern under a true selection event (bottom). Spectra represent the mean frequencies of each distinct haplotype across 10^3^ simulated replicates in a sample of *n* = 100 diploids under a constant-size simulated demographic history. Selective sweeps were simulated as one or more strongly-selected (*s* = 0.1) haplotypes rising to high frequency starting at the time of selection *t* = 400 generations before sampling.

To begin, we must first define the haplotype spectrum on which we will base our neutral expectation. That is, the spectrum that we will assign as representative of a spectrum for a genomic window under neutrality. For all genomic windows in the sample, we extract the haplotype frequency spectrum, arrange frequencies in descending order, and truncate the spectrum at an arbitrary value *K* most frequent haplotypes (compare top and middle panels of Figure 1, first column). Thus, for each window *ℓ, ℓ* = 1, 2,…, *L* for *L* windows, we have a truncated spectrum 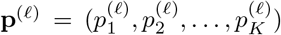, where 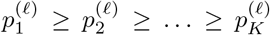, and normalized such that 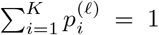. Next, we define the vector **p** = (*p*_1_, *p*_2_,…, *p_K_*), such that 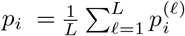 for *i* =1, 2,…, *K*. We now use **p** as our neutral expectation for likelihood computations.

From the vector **p**, we define the vector 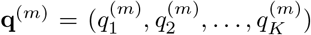, which represents a hypothetical distorted frequency spectrum consistent with a model of m sweeping haplotypes in an analysis window, with 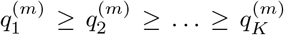. We note that our approach is purely statistical and does not feature an underlying population genetic model. We generate **q**^(*m*)^ by increasing the frequency of sweeping haplotype classes 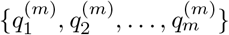 at the expense of non-sweeping haplotype classes 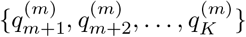. The vector **q**^(*m*)^ is related to **p** by

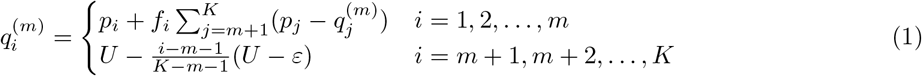

where *f_i_*, with 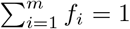 and *f_i_* ≥ 0 for *i* = 1, 2,…, *m*, is a term defining the manner in which the mass associated with haplotype frequencies {*p*_*m*+1_, *p*_*m*+2_,…, *p_K_*} in the neutral frequency spectrum is distributed among {*p*_1_, *p*_2_,…, *p_m_*} to generate the sweep frequency spectrum of the alternative model, and *U* and *ε* are respectively the frequencies of the most and least frequent non-sweeping haplotype classes, 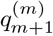 and 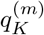.

We can define *f_i_* in multiple ways. Choosing a model of *f_i_* = 1/*m* generates an alternative model in which value is uniformly added to each of *p*_1_,…, *p_m_*. We can also specify a distortion in which value is added proportionally to each sweeping haplotype frequency, where *f*_1_ > *f*_2_ > ⋯ > *f_m_*, such as 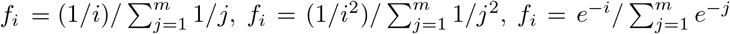, or 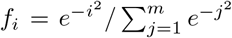. The choices of *U* and *ε* determine the frequency of the non-sweeping haplotype classes in the alternative model. For *U* > *ε*, the value of 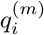 decreases linearly for *i* = *m* + 1, *m* + 2,…, *K*, whereas *U* = *ε* constrains all 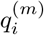 to equal *ε* for *i* = *m* + 1, *m* + 2,…, *K*. Regardless of the choice of *U* and *ε*, their relationship with each other and *p*_*m*+1_ is necessarily *p*_*m*+1_ ≥ *U* ≥ *ε*. We also note that **q**^(*K*)^ = **p** by definition, illustrating that the null (neutral) model is nested within the alternative (sweep distortion) model.

For each analysis window, we must finally obtain a vector of counts **x**, observed for the most frequent *K* haplotypes. We define **x** = (*x*_1_, *x*_2_,…, *x_K_*), where elements are once again arranged in descending order, with *x*_1_ ≥ *x*_2_ ≥ … ≥ *x_K_*. We normalize each *x_i_* to satisfy the constraint 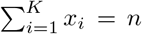, where *n* is the number of haplotypes in the sample.

Using the model haplotype frequency spectra **p** and **q**^(*m*)^ in conjunction with the observed vector of counts x for the most-frequent *K* haplotypes in a particular genomic window, we define likelihood functions, which are based on the multinomial distribution. The likelihood of the model parameters under the null hypothesis (neutrality) given the haplotype counts in an analysis window, equivalent to the probability of obtaining the observed haplotype counts **x** given **p** and *K*, is

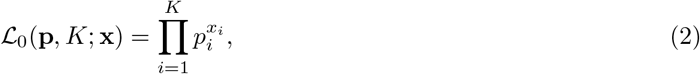

whereas the likelihood under the alternative hypothesis (sweep distortion) is

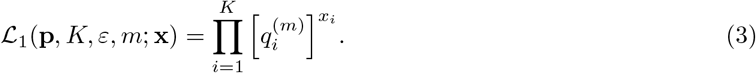

Therefore, the log-likelihoods are

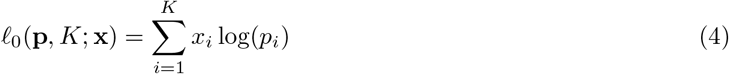

and

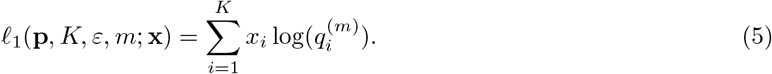

We optimize *ℓ*_1_(**p**, *K, ε, m; x*) over *m* ∈ {1, 2,…, *K*} and *ε* ∈ [1/(100*K*), *U*], keeping *U* fixed, to find

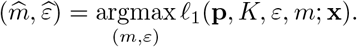

Thus, our test statistic is defined as

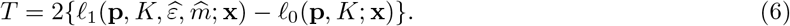

Each analysis window in the genome is assigned a test statistic in this manner, and larger test statistics indicate greater support for a sweep in the window (*i.e*., greater distortion toward few moderate- or high-frequency haplotypes). Because in the process we also identify the most likely number of presently sweeping haplotypes 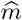 to yield the underlying distorted haplotype spectrum, our approach can also be used to quantify the softness of an identified sweep.

## Results

We first performed experiments with simulated data in which we generated populations based on nonequilibrium human demographic models [Terhorst et al., 2017], covering a variety of neutral and selection scenarios. These demographic models consisted of a history resembling that of the CEU European population, featuring a prominent bottleneck about 2000 generations prior to sampling, and a sub-Saharan African history resembling that of the YRI population, characterized by relative population effective size stability preceding an expansion. We additionally probed the effect of background selection on the value of the *T* statistic, as background selection has been implicated as a confounding factor when searching for selective sweeps [Charlesworth et al., 1993, 1995, Seger et al., 2010, Nicolaisen and Desai, 2013, Cutter and Payseur, 2013, Huber et al., 2016]. We evaluated the performance of our method in terms of ability to detect sweeps (its power) and ability to infer the number of sweeping haplotypes at the time of sampling 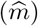, which serves as a proxy for the number of distinct haplotypes (*ν*) involved in the history of the sweep (its classification ability). Finally, we applied our method to data from the 1000 Genomes Project [Auton et al., 2015] and the *Drosophila* Genetic Reference Panel [DGRP; Mackay et al., 2012] to measure our ability to properly identify and classify selective sweep candidates.

### Detection and characterization of selective sweeps

We measured the power of our likelihood ratio test statistic (*T*) to differentiate selective sweeps from neutrality. Larger values of the *T* statistic for an analysis window indicate a greater departure from the neutral haplotype frequency spectrum and therefore a greater probability of a sweep within that genomic region. To measure power, we first simulated 1000 neutral replicates of 500 kb chromosomes under the CEU and YRI demographic models. From these simulations, we obtained each model’s expected truncated neutral haplotype frequency spectrum **p** = (*p*_1_,*p*_2_,…, *p_K_*), which was the basis of our likelihood computations (see *Theory*). The spectrum p for a model represented the mean across all genomic windows of all replicates, truncated at a particular value of *K*. Thus, *K* = 20 indicates the spectrum of the most frequent 20 haplotypes in a genomic window, whose frequencies are labeled *p*_1_ through *p*_20_. To assess power, we computed the *T* statistic for each genomic window of each simulated neutral replicate. We solely retained the maximum value of the *T* statistic across all windows for each neutral replicate, and similarly retained the maximum *T* statistic across each replicate of each selection scenario we tested. In our experiments, we assessed power at the 1 and 5% false positive rates (FPRs), meaning that we measured the proportion of selection replicates respectively exceeding the top 1 or 5% of *T* statistics within the neutral distribution.

The *T* statistic has high power to detect a hard sweep (*ν* =1 sweeping haplotype) affecting the CEU-based demographic history, regardless of selection coefficient *s* (Figure 2, top). At both the 1 and 5% FPRs, the *T* statistic reliably detects hard sweeps beginning between 1000 and 1500 generations before sampling, with stronger sweeps extending the lower bound of this range to 200 generations (Figure 2, top-right). The power of the *T* statistic attenuates for more ancient sweep events because haplotype identity surrounding the selected site decays over time in the population as mutation and recombination generate new haplotypes. Additionally, power to detect the most recent weak sweeps is low because sufficient time has not elapsed for the selected haplotype to reach high frequency. The *T* statistic achieves greater power for simulated YRI demographic models than for CEU models across all tested scenarios (Figure 2, bottom). This increased power is due to the greater effective size of African relative to European human populations, which results in greater background haplotype diversity and therefore increased prominence of selective sweeps. Accordingly, power declines more slowly for older sweeps, and remains for sweeps as old as 4000 generations before sampling. Choosing alternate values of *K* (10, 15, or 25; Figure S1) yielded little change in power to detect simulated sweeps from *s* ∈ [0.005,0.5] relative to *K* = 20 (Figures 2, middle column) at the 1% FPR, with power at the 5% FPR slightly larger for smaller values of *K*.

**Figure 2:**
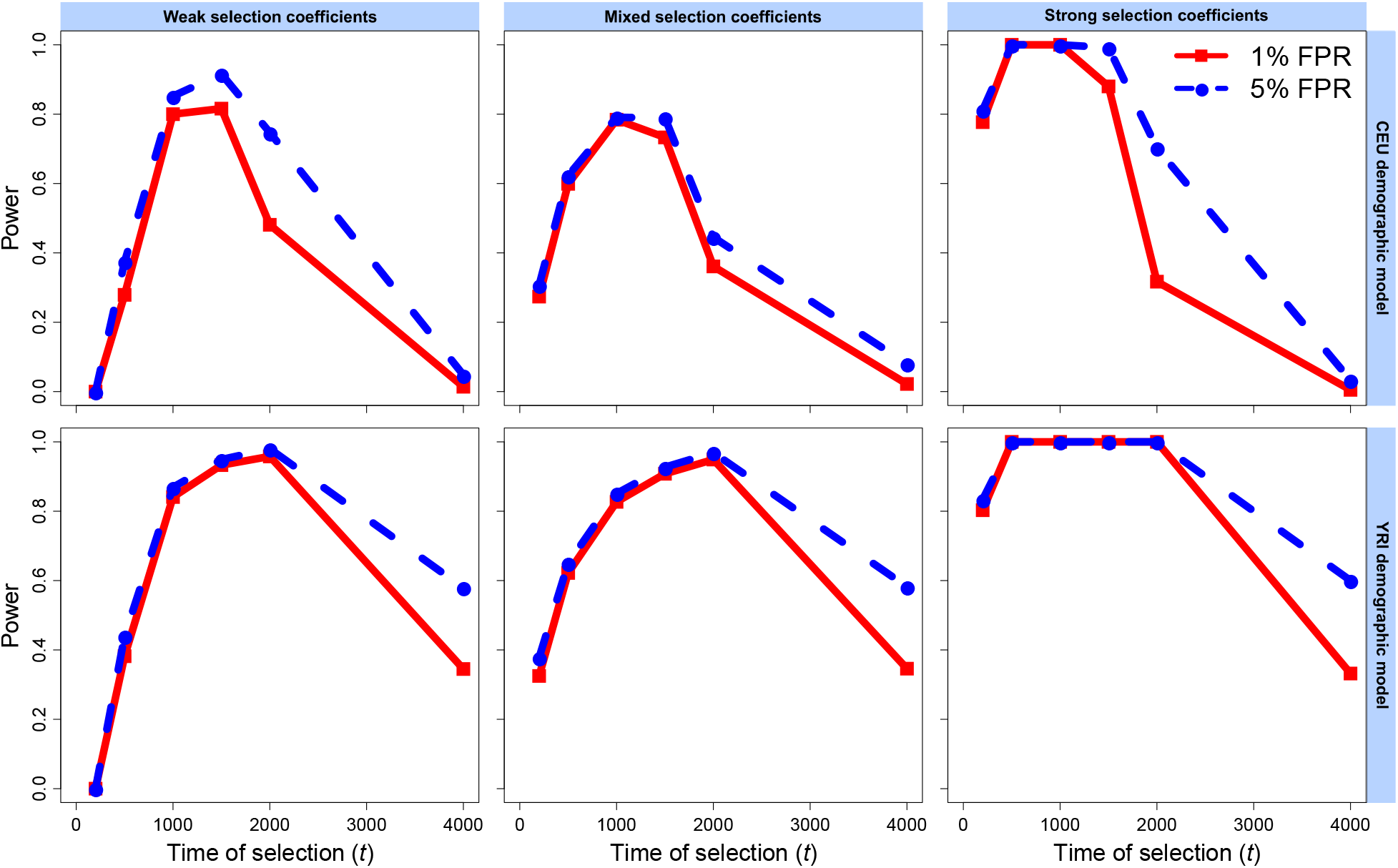
Power of the *T* statistic at 1% and 5% false positive rates (FPRs) to detect hard selective sweeps from a single *de novo* mutation arising at time *t* ∈ {200, 500,1000,1500, 2000, 4000} generations before sampling under the European CEU (top) and sub-Saharan African YRI (bottom) human demographic models, for phased haplotypes. Weak selection coefficients were drawn uniformly at random from *s* ∈ [0.005, 0.05], strong selection coefficients were drawn uniformly at random from *s* ∈ [0.05, 0.5], and mixed selection coefficients were drawn uniformly at random on a log-scale from *s* ∈ [0.005, 0.5]. All inferences used a spectrum of *K* = 20 for likelihood computations.

For soft sweeps from selection on standing genetic variation (SSV, *ν* ∈ {2, 4, 8,16, 32}; Figure 3), the power of the *T* statistic attenuates more rapidly than for hard sweeps, and *T* never reaches values as large, especially for weaker sweeps. Under both CEU (Figure 3, top) and YRI (Figure 3, bottom) demographic histories, trends in power remain similar regardless of the number of sweeping haplotypes, with maximum power of *T* achieved for sweeps between 1000 and 1500 generations old; however, power declines as the number of sweeping haplotypes increases. Assessing power at the 5% FPR indicates that we nonetheless maintain extensive differentiation between sweeps and neutrality for up to *ν* = 8 distinct initially sweeping haplotypes drawn from selection coefficients *s* ∈ [0.005, 0.5] for CEU models, or up to *ν* = 16 for YRI models (mixed sweeps). This remains true for weaker (*s* ∈ [0.005,0.05]) and stronger (*s* ∈ [0.05, 0.5]) sweep sets under the CEU model, but for YRI we see reliable power for 16 sweeping haplotypes at the 1% FPR across all sweep strengths, and for weak sweeps we retain high power at the 5% FPR for 32 sweeping haplotypes. Thus, the demographic history of the sampled population plays an important role in the power of the *T* statistic, consistent with the results of other haplotype-based approaches [Harris et al., 2018a], but our results indicate that the *T* statistic is nonetheless flexible as to the selection scenarios that it can distinguish from neutrality.

**Figure 3:**
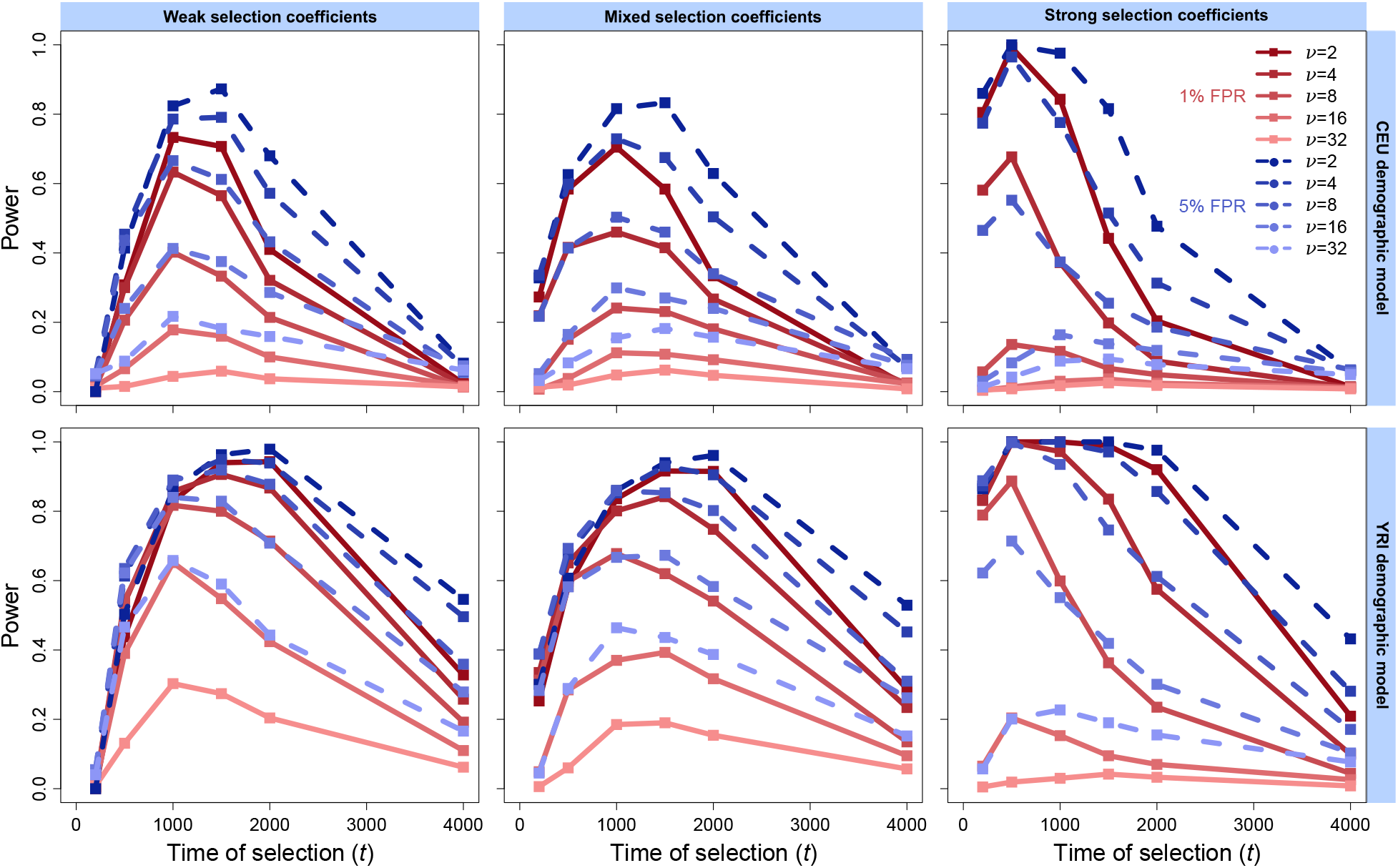
Power of the *T* statistic at 1% and 5% false positive rates (FPRs) to detect soft selective sweeps from selection on standing variation on *ν* ∈ {2, 4, 8,16, 32} distinct sweeping haplotypes beginning at time *t* ∈ {200, 500,1000,1500, 2000, 4000} generations before sampling under the European CEU (top) and sub-Saharan African YRI (bottom) human demographic models, for phased haplotypes. Weak selection coefficients were drawn uniformly at random from *s* ∈ [0.005, 0.05], strong selection coefficients were drawn uniformly at random from *s* ∈ [0.05, 0.5], and mixed selection coefficients were drawn uniformly at random on a log-scale from *s* ∈ [0.005, 0.5]. All inferences used a spectrum of *K* = 20 for likelihood computations.

Selective sweeps produce elevated values of the *T* statistic along the simulated chromosome that on average peaks in the region surrounding the site under selection (Figures S2 and S3, first and third rows). Furthermore, *T* remains elevated beyond the 450 kb bounds that we examined, indicating that on average, the shape of its distribution in a genomic region, as well as its overall elevated value, can be used to distinguish selection from neutrality under scenarios in which we have power. A signal peak remains even for scenarios in which we do not have high power, though its maximum associated value remains small on average. Because neutral regions are likely to feature plateaus rather than peaks in the value of the *T* statistic, our observations illustrate the potential importance of considering the correlation in signal between windows to identify more subtle selection signatures. This is especially important for soft sweeps, which lose prominence proportionally to the number of sweeping haplotypes, but still produce a peak-like distortion of local *T* statistic values.

In addition to evaluating the power of the *T* statistic, we measured the ability of our approach to infer the number of presently sweeping haplotypes 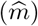 at the site under selection. The ability to infer 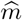 is a result of optimizing the likelihood function *ℓ*_1_ over all possible *m* for the chosen truncation *K* (see *Theory*). In Figure 4, we show the distribution of *T* statistics with their associated haplotype frequency spectra and 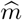, for each of 10^3^ neutral, mixed hard sweep (*s* ∈ [0.005, 0.5], *ν* =1, *t* = 1000), and mixed soft sweep (*ν* = 4) replicates, under both the CEU and YRI models (same data as Figures 2 and 3). Relative to neutrality (Figure 4, left), we more often assign smaller 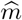 (indicated by black and darker purples) to sweep simulations (Figure 4, center and right). This result fits with the expectation that under a sweep, the first few haplotype classes exist at elevated frequency relative to the remaining classes, and this also translates to larger values of *T* for those replicates. Accordingly, sweeps that are weaker due to their age or selection coefficient are not only difficult to detect, but also difficult to classify, yielding patterns that fit within the neutral distribution. We found that trends were highly congruent between the CEU and YRI sweep models, but the large neutral background diversity for YRI made it less likely that we would infer a small 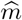 in the absence of a sweep.

**Figure 4:**
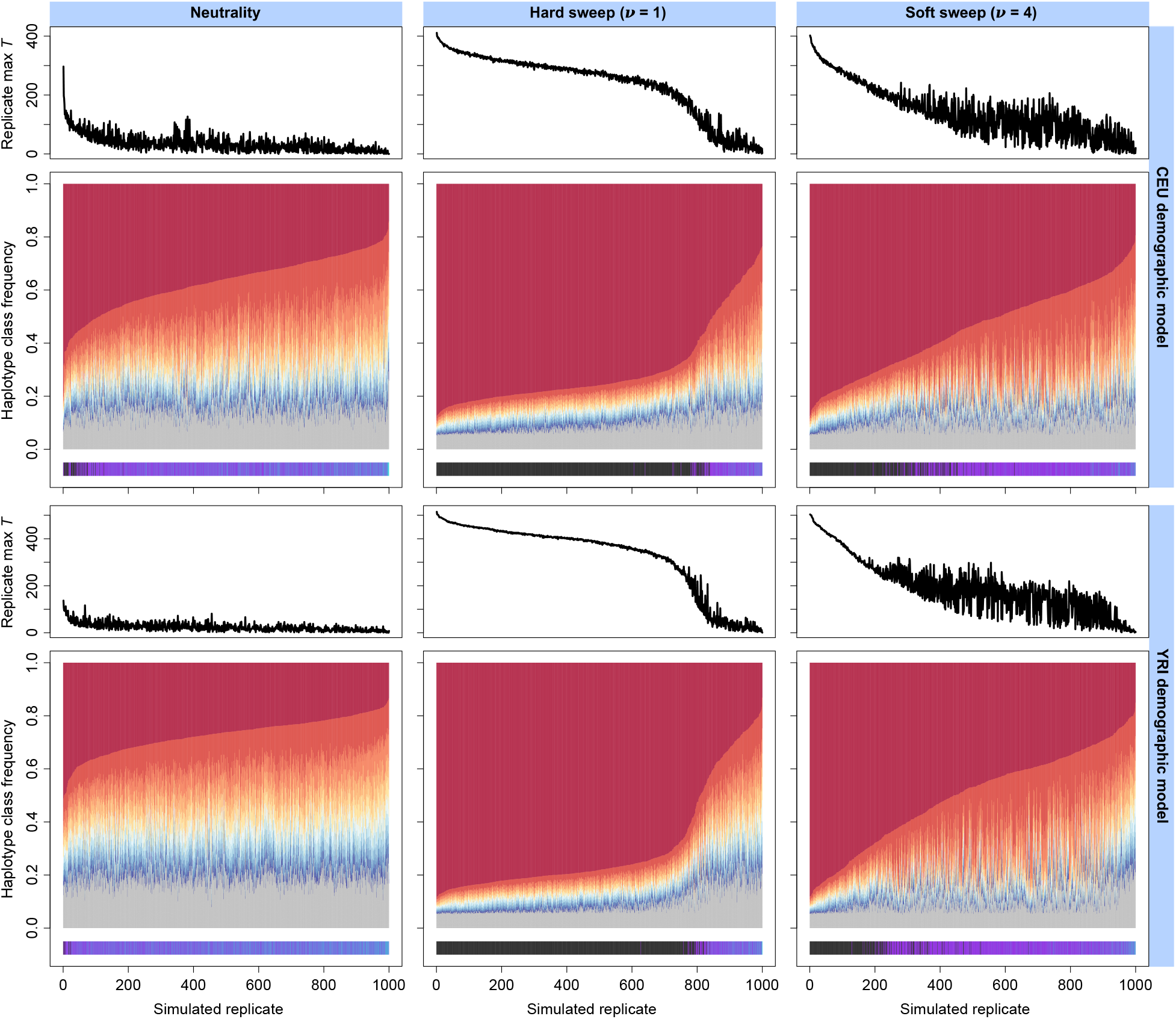
Truncated haplotype frequency spectra (*K* = 20) across 10^3^ simulated replicates for analysis window of maximum replicate-wide *T* statistic under neutral (left), hard sweep (center), and soft sweep (right) scenarios, for European CEU (top) and sub-Saharan African YRI (bottom) human demographic models. Each simulated replicate is one vertical slice within the greater plot, and the 10 most frequent haplotypes are colored on a scale from red (most-frequent) to blue (10th most-frequent), while the remaining haplotypes are shaded together in gray. Replicates are associated with their *T* statistic (above) and their inferred 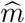 (below). Inferred hard sweeps 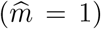 are indicated in black, whereas inferred soft sweeps 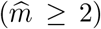 are indicated on a color scale spanning purple (fewer sweeping haplotypes) to teal (maximum of 20 sweeping haplotypes, consistent with neutrality). Replicate spectra are arranged in decreasing order of most-frequent haplotype frequency.

Expanding upon Figure 4, we generated box plots summarizing the distribution of 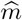 across each mixed-strength sweep scenario we previously analyzed (Figures 2, 3, and S1). In this way, we were able to better understand our ability to correctly classify sweeps as hard or soft, as well as understand the relationship between the initialized number of sweeping haplotypes within our simulations and the observed number of sweeping haplotypes at the time of sampling. The most accurate inferences of 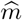 with respect to *ν* overlapped the scenarios in which the *T* statistic had the greatest power. This comprised selective sweeps beginning between 500 and 2000 generations before the time of sampling (Figures S4-S6). For hard sweeps under either demographic model, we were able to consistently infer a median of one sweeping haplotype from the genomic window of maximum *T* within *t* ∈ [500, 2000], losing accuracy outside of this range (Figures S4-S6). Soft sweep scenarios invariably displayed an increase in 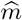 relative to hard sweeps that corresponded with increases in the number of initially-selected haplotypes (*ν*), but at the time of sampling, 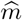 was typically considerably smaller than *ν* (Figures S4 and S5). Furthermore, older soft sweeps, up to 2000 generations old, were associated with fewer sweeping haplotypes than younger sweeps for all *ν*, indicating the loss of particular sweeping haplotypes over the course of selection, and prior to the decay of the overall sweep signature in the sample. For all scenarios, we also found that the mean value of 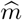 along the simulated chromosome followed an inverse trend to the mean value of the *T* statistic, forming a valley where the *T* statistic forms a peak (Figures S2-S3).

Because phasing haplotypes may not be possible in all cases, such as in the study of non-model organisms, we sought to expand our application of the *T* statistic to unphased multilocus genotype (MLG) data. To evaluate power for MLGs, we reused the previous simulated human demographic model replicates of prior experiments (represented in Figures 2 and 3), merging each individual’s two haplotypes. Whereas haplotypes are character strings indicating the state of a biallelic SNP as either reference or alternate along a region of one copy of an individual’s genome, MLGs have three possible states for each biallelic SNP—homozygous reference, homozygous alternate, or heterozygous—and half the sample size of phased haplotypes. We found that, as with the transition between phased and unphased data for haplotype homozygosity statistics [Harris et al., 2018a, Harris and DeGiorgio, 2019], power for the unphased application of the *T* statistic was wholly consistent with that of the phased application, for both hard (Figure S7) and soft (Figure S8) sweeps. As we expected, the smaller size of the MLG samples resulted in slight decreases in power for each sweep scenario, as well as smaller values of the *T* statistic relative to the phased application, but our results indicate that selective sweeps may be reliably identified nonetheless without the need to phase haplotypes. Likewise, we found that the *T* statistic applied to MLGs could generate inferences of 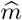 that matched those of haplotype data, further underscoring the parallel performance of our approach on unphased data (Figure S9).

We examined background selection scenarios for both haplotype and MLG data to determine whether the loss of genetic diversity associated with linked purifying selection could spuriously yield elevated values of the *T* statistic. Simulating 500 kb chromosomes as previously under both human demographic models, we found that background selection had no effect on the distribution of *T* relative to neutrality. We determined this by observing the receiver operating characteristic curves comparing neutral scenarios to those in which a central gene experiences strong (*s* = −0.1) background selection for the duration of the simulation (see *Materials and Methods*). For both the CEU (Figure S10, top) and YRI (Figure S10, bottom) populations, across central genes of size 11 kb (Figure S10, left) and 55 kb (Figure S10, right), we see that all curves fit tightly along the diagonal, indicating no difference between compared replicate sets. Therefore, we expect that the presence of background selection, for which we do not explicitly account in our model, should not affect inferences with the *T* statistic.

### Application to empirical datasets

We searched for candidate selective sweeps in human and *D. melanogaster* datasets using the *T* statistic, choosing these datasets because of their high quality, size, and availability of phased haplotypes. Specifically, the 1000 Genomes [Auton et al., 2015] dataset contains no missing data, as all allelic states have been imputed. Meanwhile, the DGRP [Mackay et al., 2012] dataset provides a classic invertebrate model whose properties deviate considerably in history and genomic architecture from the mammalian model of humans. We obtained values of *T* and 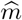 for each genomic window, and assigned to each gene a *T* and 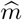, as well as a *p*-value, based on the window of maximum *T* overlapping that gene. For a window to be associated with a gene, its central SNP must lie between the transcription start and stop sites of the gene. For human analyses, we scanned with a 117-SNP window advancing at increments of 12 SNPs, and for *D. melanogaster*, we used windows of size 400 SNPs advancing by 40 SNPs, as in Garud et al. [2015] and Harris et al. [2018b], to which we compare our results. Both window sizes were based on the minimum window size across which LD had decayed beyond one-third of the LD between SNPs separated by one kb in order to eliminate the effect of background LD on inferences (see *Materials and Methods*). We also analyzed the human dataset as MLGs by manually merging an individual’s two haplotypes together within a window, allowing us to demonstrate the performance of the *T* statistic following its successful application to unphased simulated data (Figures S7 and S8). We did not need to distinguish between haplotype and MLG approaches for the *D. melanogaster* dataset because the study population consisted of only inbred individuals, rendering the distinction between phased and unphased data meaningless.

For human data, we examined the CEU and YRI populations (Tables S1 and S2), matching the demographic models used in our simulations. Among the top 40 sweep candidates of either population, hard sweeps predominated within the phased haplotype data, comprising all but two top candidates among the CEU, and 67.5% of top candidates among the YRI. Additionally, each of these candidate soft sweeps, except for *BTNL2* in YRI 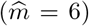 featured only three or fewer sweeping haplotypes. This result indicates that the *T* statistic is more sensitive to harder sweeps than to softer ones, which is a consequence of the greater distortion in the haplotype frequency spectrum of hard sweeps relative to soft sweeps. This finding matches our simulated results, in which the value of *T* was proportional to the number of sweeping haplotypes in the population. Moreover, the increased presence of candidate soft sweeps in YRI relative to CEU mirrors our observation from simulated data that the *T* statistic has greater power to detect softer sweeps for populations that have not experienced a bottleneck in their history. Furthermore, these patterns corroborate results from the H12 analysis of this dataset [Harris et al., 2018a], which found more hard sweeps than soft in the CEU population, and among top candidates generally.

Across both the CEU and YRI populations, we were able to recover most top candidates from the haplotype data within the MLG data, indicating the reliability of using MLGs for inference with the *T* statistic in natural populations when phased data are unavailable. The MLG results deviated somewhat from the haplotype results, however, when classifying candidates as hard or soft. Multiple candidates inferred to be hard sweeps from the haplotype data were classified as soft from their MLG spectra. These candidates include *XIRP2* and *BCAS3* in the CEU population, as well as *ITGAE, SUGCT, NNT*, and *HLA-DPB2* in the YRI population. We examine the latter candidate more closely in Figure 5. These differing inferences may arise from the slightly different interpretation of 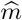 between phased and unphased data. For phased data, 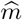 refers to the number of sweeping haplotypes, whereas for unphased, it measures the number of sweeping MLGs, which may be different for the same genomic window between the different data types. We also note that multiple top candidates in the MLG data inferred as soft are simply not present among top haplotype candidates, indicated by the absence of a turquoise-colored background in Tables S1 and S2. We consider the application of the *T* statistic to MLGs further in the *Discussion*.

**Figure 5:**
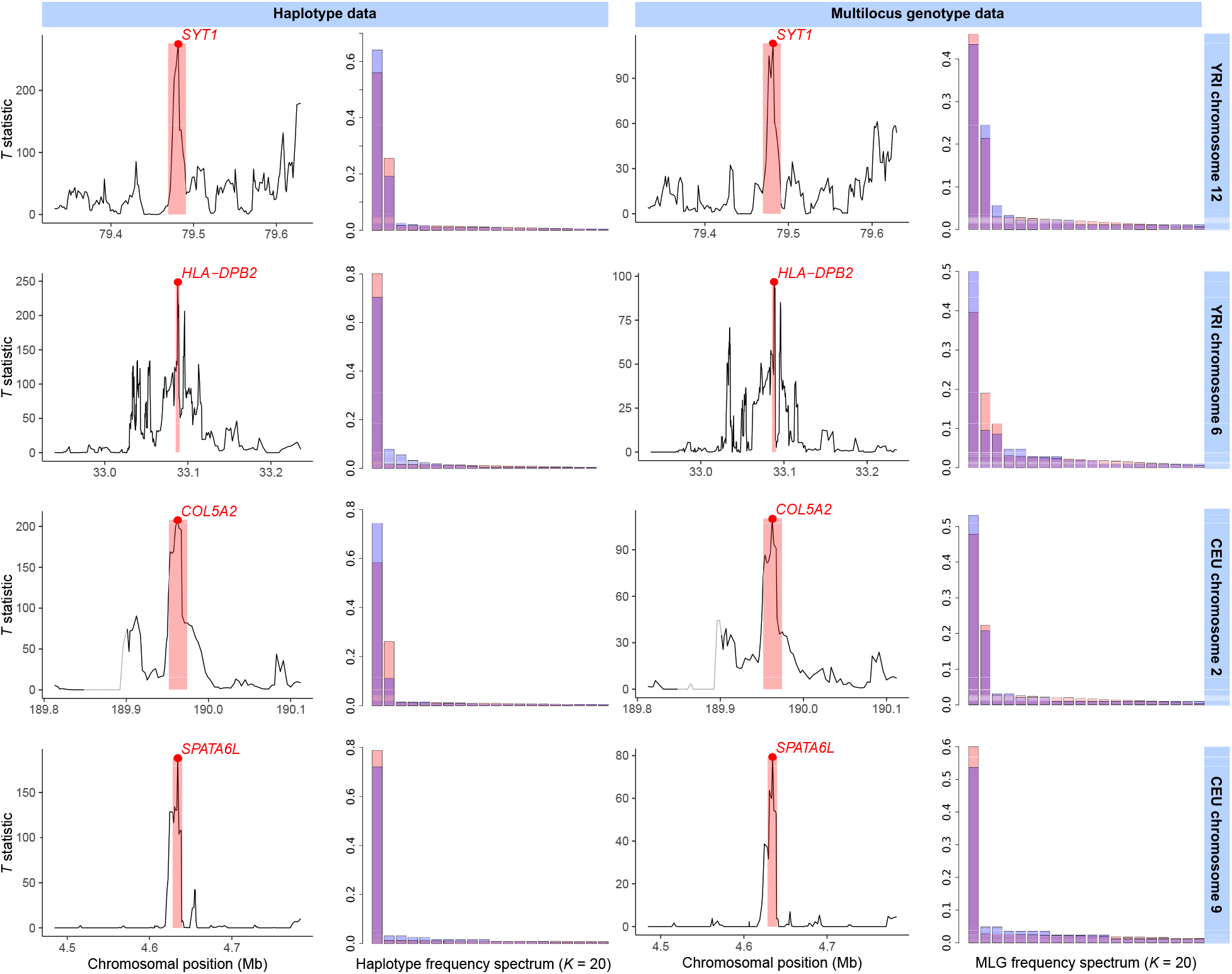
Selective sweep candidates detected with the *T* statistic from the 1000 Genomes Project dataset [Auton et al., 2015] as phased haplotypes (left) and unphased multilocus genotypes (MLGs, right). For each of four sweep candidates in the human YRI (top two rows) and CEU (bottom two rows) populations, we show the *T* statistic across the 300 kb interval surrounding the candidate peak, as well as the frequency spectra for the most likely sweep model corresponding to the candidate at the 117-SNP analysis window of maximum *T*. The window of maximum *T* is shaded in red, with the position of the window center (median SNP) as a red dot. The frequency spectrum of the most likely model is also shown in red, whereas the observed frequency spectrum at the point of maximum *T* is overlaid in blue. The displayed candidates are a putative soft sweep 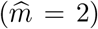 at *SYT1* in YRI (top row), hard sweep 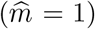 at *HLA-DBP2* in YRI (second row), soft sweep 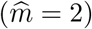 at *COL5A2* in CEU (third row), and hard sweep at *SPATA6L* in CEU (bottom row). The gray segment upstream of *COL5A2* (third row) indicates a portion of the genome that was filtered out (see *Materials and Methods*).

Among top sweep candidates in human data were expected results, including a hard sweep 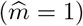 at the cluster of genes on CEU chromosome 2 comprising *LCT, MCM6, DARS*, and *ZRANB3* (minimum *p*-value < 10^−6^), related to a well-documented adaptation to milk-based diets in European populations [Bersaglieri et al., 2004]. Additionally, we found two top candidates in CEU that have not been explicitly described as sweeps previously, *RSPH3* and *ZNF211* (both 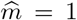). *RSPH3* encodes a radial spoke protein that is integral in the structure of 9 + 2 motile cilia across diverse cell types, including flagellated cells [Teves et al., 2016], and so selection here may be related to ancient sperm competition in humans [Leivers et al., 2014]. *ZNF211* is among a diverse set of zinc-finger genes whose products are believed to participate in the inactivation of endogenous retroviruses, parasitic mobile DNA whose effects can be deleterious to their hosts [Lukic et al., 2014]. We recovered *SYT1* (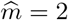; *p* = 10^−6^), *NNT* 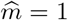, *HEMGN* 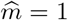, and *RGS18* 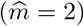 in YRI, which have all received attention as potential adaptive targets [Voight et al., 2006, Pickrell et al., 2009, Fagny et al., 2014, Harris et al., 2018a]. In addition, both populations yielded *HLA* genes as top sweep candidates, overlapping at *HLA-DRB5* 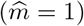, whereas *HLA-DPB1* 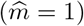 was exclusive to CEU and *HLA-DPB2* (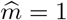 *p* = 2 × 10^−6^) was exclusive to YRI. This shared signal supports the recent evidence [Albrechtsen et al., 2010, Goeury et al., 2017] that sweeps at HLA loci, including those which we describe here, were important in the development of modern genetic diversity in human immune-related genes.

In Figure 5, we take a closer look at top candidate hard and soft sweeps uncovered in our scan of the 1000 Genomes dataset [Auton et al., 2015], across both haplotypes and MLGs. Each top candidate fell within a well-defined *T* statistic peak region surrounded by regions of low signal, and this spatial signature was consistent between both data types. First, we found *SYT1* as a significant (*p* = 10^−6^) top soft sweep in the YRI population, featuring both 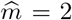 sweeping haplotypes and 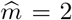 sweeping MLGs at the window of maximum signal (Figure 5, first row). *SYT1* is the cell surface receptor through which the type B botulinum neurotoxin of *Clostridium botulinum* bacteria enters human neurons [Connan et al., 2017], and so a sweep here may be involved in resistance to this infection [Harris et al., 2018a]. Next, we identified *HLA-DPB2* as a significant hard sweep in YRI (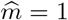; *p* = 2 × 10^−6^) based on haplotypes, but featuring three elevated MLGs within the window of maximum signal (Figure 5, second row). Looking at the haplotype frequency spectrum, it is clear that one haplotype predominates, and equivalently, only one MLG predominates, but individuals heterozygous for the first haplotype and either the second or third comprise just under 20% of the population, leading to an inference of 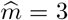. *COL5A2* was the most outlying soft sweep candidate we identified in CEU, harboring 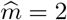 inferred sweeping haplotypes, but with a sevenfold disparity between their frequencies. This gene has received little attention, but is located within a significantly overrepresented run of homozygosity associated with schizophrenia [Lencz et al., 2007]. Finally, we propose the spermatogenesis-associated protein *SPATA6L* as a hard sweep candidate in CEU. Our finding here of an isolated *T* peak fits with existing evidence of selection at other spermatogenesis proteins [Schrider and Kern, 2017], and with the result that European and sub-Saharan African populations are diverged at this locus, with selection in the hunter-gatherer Batwa population inferred here [Bergey et al., 2018].

Our scan of the North American DGRP population of *D. melanogaster* also identified expected sweep candidates among the top genic *T* statistic peaks. We note that while we were unable to establish statistical significance against a neutral model based on the DGRP demographic history of Duchen et al. [2013] (see *Materials and Methods*), our top candidates have literature support as potential adaptive targets. Foremost among functionally-characterized candidates was *Ace*, which encodes the acetylcholinesterase enzyme and has long been implicated in the development of resistance to organophosphate and carbamate insecticides within *D. melanogaster* [Menozzi et al., 2004, Karasov et al., 2010, Garud et al., 2015]. However, contrary to previous studies alleging a soft sweep at *Ace* [Karasov et al., 2010, Garud et al., 2015], we found the greatest support for a model of only one sweeping haplotype. We identified another candidate hard sweep of similar magnitude at *Uhg1*, which also contributes to insecticide resistance, but to the organochlorine DDT [Pedra et al., 2004]. The methyltransferase-encoding gene *Pimet* emerged as the most prominent candidate soft sweep 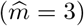 in our search, and is central to the viral RNA degradation pathway that is subject to ongoing coevolution against pathogen incursion and deleterious transposable element activity [Kolaczkowski et al., 2011, Lee and Langley, 2012]. We finally highlight *ana3* as a candidate for adaptation in *D. melanogaster*. This prospective hard sweep affects a highly-conserved gene encoding a centriole protein fundamental to the structural integrity of basal bodies within cells [Stevens et al., 2009]. A sweep here may contribute to enhanced success in sperm competition, and fits with the expectation that sperm gene evolution is an ongoing and central part of positive selection in *D. melanogaster* [Nurminsky et al., 1998, Dorus et al., 2008, Wong et al., 2008, Yeh et al., 2012].

## Discussion

We have proposed a likelihood-based approach to detect selective sweeps in whole-genome polymorphism data that is applicable to a variety of different demographic scenarios, classifies detected sweeps as hard or soft without needing to rely on additional analyses or statistics, and is the first likelihood-based method to leverage distortions in the haplotype frequency spectrum to make inferences. Each of these attributes is important because selective sweeps are multifaceted genomic signatures that are not always characterized by the presence of few high-frequency haplotypes [Jones et al., 2013, Wilson et al., 2017], may be ongoing or incomplete at the time of sampling [Vy and Kim, 2015, Vy et al., 2017], and may range in strength across multiple orders of magnitude [Messer and Neher, 2012, Nam et al., 2017]. Thus, our simulation experiments probed a realistically diverse complement of sweep scenarios likely to be relevant to a variety of study systems. Most importantly, the *T* statistic demonstrated high and consistent power and classification ability across examined parameters, highlighting its suitability to make inferences within variable contexts.

Expectedly, the *T* statistic had the greatest power to identify recent selective sweeps on fewer haplotypes, and lost power proportional to the extent of departure from these ideal conditions (Figures 2 and 3). Because it is haplotype-based, the *T* statistic captures distortions in the haplotype frequency spectrum relative to neutral expectations. These distortions require time to establish, and decay over time as well. Thus, we found that for human demographic models, the *T* statistic could reliably identify sweeps occurring within 2000 generations of sampling. For stronger sweeps, power was consistently elevated across this range, but because weaker sweeps require more time to establish, this range narrows as sweep strength decreases. Additionally, we uniformly had more power to detect sweeps under the YRI demographic model than the CEU. This is due to the severe bottleneck underlying the history of the CEU, as well as all non-African human populations. Bottlenecks may reduce the diversity of haplotypes within a population, reducing the distinctiveness of sweeps relative to neutrality, whereas population expansions have the opposite effect [Jensen et al., 2005, Campbell and Tishkoff, 2008]. Nonetheless, we could generally detect sweep strengths across orders of magnitude between 1000 and 2000 generations before sampling under either demographic model, comprising selective events that in humans cover the period from 25,000 to 58,000 years ago, between the out-of-Africa event and the spread of agriculture [Nakagome et al., 2015].

The choice of human demographic history did not impact our inference on the number of currently sweeping haplotypes 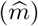 in the population, and we found that inferred 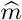 provided a generally accurate representation of sweep history that distinguished between hard and soft sweeps (Figures 4 and S4-S6). Across all experiments, we found that as long as the *T* statistic has power to detect a sweep, it would assign an appropriate 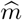, and often continue to do so even after power had waned, especially under the CEU history (Figures S4-S6). Accordingly, we observe a distortion in the spatial signal of 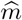 mirroring that of *T*, which can remain distorted surrounding the site under selection for sweeps older than 2000 generations (Figures S2 and S3). Among detectable sweeps, approximately 80% of mixed strength hard sweeps (*s* ∈ [0.005,0.5], *ν* =1) beginning at *t* = 1000 generations before sampling were identified as hard 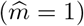 as long as they established in the population (Figure 4, middle), whereas more than 60% of mixed strength soft sweeps (*ν* = 4) yielded 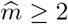 (Figure 4, right). Because haplotypes may be lost over the course of selection due to drift, we limit our inferences to the binary choice of hard or soft, and we acknowledge that 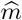 is likely to be a better proxy for *ν* in cases of fewer sweeping haplotypes, as most sweeping haplotypes are likely to be lost for sweeps from larger *ν* (Figures S2 and S3).

As an attempt to improve the performance of the *T* statistic, We sought to examine whether the choice of sweep distortion model, based on the choice of *f_i_* (see *Theory*), would affect our inferences. Ultimately, we found that all of our tested models yielded little difference in the power of *T* to identify sweeps (Figure S12). The five models we examined, consisting of (A) 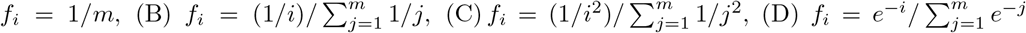, and 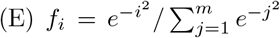, differ in the amount of
 weight allocated to the secondary sweeping haplotypes 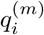 for *i* ∈ {2, 3,…, *m*} relative to 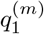 when distorting **p**. In model A, which we use as our default approach, each sweeping haplotype gains the same amount of weight after distortion, ensuring that each is prominent within spectrum **q**^(*m*)^. Models B through E represent increasingly uneven weight distributions that favor frequency 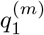 at the expense of 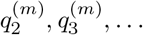, and 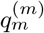, which we proposed based on our observation in simulated data that soft sweeps do not affect each sweeping haplotype evenly, and one sweeping haplotype may still be considerably more prominent than the rest (Figure 1). Furthermore, the different *T* statistic variants demonstrated little difference in sweep classification ability (Figure S13), suggesting that the most important consideration in constructing our sweep models is simply defining the number of sweeping haplotypes, and not the manner in which they sweep.

An important feature of the *T* statistic is its ability to detect sweeps from unphased MLG data. Our ability to extend the power of our approach to MLGs is meaningful because it provides the ability to interrogate polymorphism data from non-model organisms for which phased haplotypes are unavailable, difficult to obtain, or unreliable [Browning and Browning, 2011, O’Connell et al., 2014, Castel et al., 2016, Laver et al., 2016, Zhang et al., 2017, Harris et al., 2018a]. Overall, we found no difference in power trends between the two data types, such that scenarios under which we have high power with phased data are scenarios of high power with unphased data (compare Figures 2 and 3 to Figures S7 and S8). However, we find that the *T* statistic applied to haplotypes always matched or exceeded power for MLG data. This is to be expected because MLGs are a more diverse data type. Under a random mating assumption, the presence of a single high-frequency haplotype implies that only one MLG should exist at high frequency, but in the case of two high frequency haplotypes, both homozygotes, as well as their heterozygote, will be prominent in the population. In this way, a sweep on two haplotypes can appear as a sweep on three MLGs, and sweeps on larger numbers of haplotypes will result in even larger numbers of elevated MLGs, which may be more difficult to separate from neutrality. Likewise, one high frequency haplotype and one medium frequency haplotype can yield two high frequency MLGs, meaning that an inferred 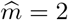 in MLG data could underlie a true hard sweep. Fortunately, our results indicate that sweep classification across phased and unphased datasets is consistent in simulated data (Figure S9), and largely so in practice (Tables S1 and S2).

We further contextualized the power of the *T* statistic by directly comparing it to that of the H12 and G123 statistics for phased [Garud et al., 2015] and unphased [Harris et al., 2018a] data, respectively (Figures S14-S17). H12 and G123, which we will collectively call the identity statistics, are expected homozygosity measures equaling the sum of squared haplotype (H12) or MLG (G123) frequencies wherein the two (H12) or three (G123) largest frequencies are pooled together into one. Increased levels of observed homozygosity are a classic signature of selective sweeps [Sabeti et al., 2002], whereas pooling provides an increased power to identify soft in addition to hard sweeps [Garud et al., 2015]. Applying the identity statistics to the same simulated human-modeled data we analyzed previously with the *T* statistic, and using the same 117-SNP window size, we found that the *T* statistic had a modest but consistent advantage in power over the identity statistics, which was particularly apparent for the CEU model (compare Figures S14-S17 to Figures 2, 3, S7, and S8). The power of the identity statistics only matched that of the *T* statistic over a narrow interval for the strongest of sweeps, and the signal of the identity statistics faded considerably more rapidly than did that of the *T* statistic. Consequently, we expect that our likelihood-based approach to detect selective sweeps should be preferable to the identity statistics in most cases. Due to both approaches’ reliance on distortions in the haplotype frequency spectrum, and their usefulness for classifying sweeps as hard or soft (discussed further below), we believe that the *T* statistic represents an appropriate successor to the model-free identity statistics.

Further cementing this notion, the results from our scans of whole-genome polymorphism datasets (Tables S1-S3) primarily corroborated prior analyses with H12 and G123 [Garud et al., 2015, Harris et al., 2018a], as well as others. For both phased and unphased data, our top candidates in the CEU population were centered on the *LCT* locus associated with adaptation to dairy consumption [Bersaglieri et al., 2004], while expected top candidates in YRI included *SYT1, NNT, LONP2*, and *HEMGN* [Voight et al., 2006, Pickrell et al., 2009, Fagny et al., 2014, Pierron et al., 2014]. Meanwhile, all of our top candidates in the scan of DGRP data overlapped with one of the top 10 H12 peaks identified by Garud et al. [2015], except for *Uhg1* (which was in a lower-ranked peak), *nompC*, and *jar*. The advantage of the identity statistics was that they were among the first methods to enhance our understanding of selective sweep events by classifying them as hard or soft from easily interpretable summary statistics. These summary statistics, H2/H1 and G2/G1, in conjunction with their respective sweep statistic, H12 or G123, provided a reliable classification framework that suffered from the laboriousness of determining the numerical cutoffs separating hard and soft sweeps. To determine the number of sweeping haplotypes implied by paired (H12, H2/H1) or (G123, G2/G1) values required the use of approximate Bayesian computation (ABC), in which millions of demographically realistic sweep models would need to be simulated to create a distribution of paired values to which each sweep candidate would be compared. In contrast, our present approach classifies sweeps by simply selecting the most likely sweep model through optimization over *m*, requiring no knowledge of the study population’s demographic history as it is approximated by the genome-wide haplotype frequency spectrum used for the null model.

We believe that our *T* statistic will serve as an important contribution to the field of selective sweep detection methods, providing the first maximum-likelihood approach that exploits a haplotype and MLG frequency spectrum distortion model. As such, the *T* statistic is well suited to the identification and classification of recent selective events in natural populations. Indeed, our lack of dependence on phased data provides the opportunity to search for sweep signatures in any non-model organism for which whole-genome polymorphism data exist. Meanwhile, our use of multilocus data insulates us from misidentifying background selection as a sweep, a consideration for which site frequency spectrum-based inferences have to account [Enard et al., 2014, Huber et al., 2016]. We expect that our simple yet powerful statistical model of selective sweeps will yield novel insights into the adaptive histories of diverse populations, and given its suitability to human data, will prove important in future analyses of understudied populations. To motivate this point, we highlight that insights into local adaptation within human populations continue to emerge [Hu et al., 2017, Buckley et al., 2017, Fan et al., 2019], more than a decade after the first investigations began [Ronald and Akey, 2005, Bustamante et al., 2005, Sabeti et al., 2006]. To facilitate the adoption of our *T* statistic, we provide open-source software, titled `LASSI` (**L**ikelihood-based **A**pproach for **S**elective **S**weep **I**nference; http://personal.psu.edu/mxd60/LASSI.html), which implements all stages of our methodology in a single efficient pipeline.

## Materials and methods

We applied the *T* statistic to simulated data based on demographic models consistent with recent estimates of human [Terhorst et al., 2017] and *D. melanogaster* [Duchen et al., 2013] population history. Across all applications, we fixed model parameter *U* = *p_K_*, and optimized *ε* over the interval *ε* ∈ [1/(100K),*U*]. For some experiments evaluating power under human models, we also applied the *T* statistic to unphased multilocus genotype (MLG) data, which we produced by manually merging each simulated individual’s two haplotypes. We generated these data using the population-genetic simulation software SLiM [Haller and Messer, 2017], in conjunction with the coalescent simulator *ms* [Hudson, 2002]. For power experiments based on human models, simulations were performed forward in time exclusively with SLiM following a Wright-Fisher model [Fisher, 1930, Wright, 1931, Hartl and Clark, 2007]. These simulations lasted for a total of 200,000 generations, of which the former 100,000 (equivalent to 10*N*, where *N* = 10^4^ is the diploid effective population size of the simulated population) was a burn-in period to achieve equilibrium values of neutral variation, and the latter 100,000 was the period over which population size variation occurred. Simulations were scaled by a factor of λ = 20 to speed up run time, where mutation rates, recombination rates, and selection coefficients were multiplied by λ, while the size of the simulated population and the duration of the simulation in generations was divided by λ. Thus, simulation duration was reduced by a factor of 400.

For all other simulations, we generated data for each replicate population using *ms*. The outputs of these simulations were used directly to compute *p*-values (see below). For human simulations, we chose a mutation rate of 1.25 × 10^−8^ per site per generation [Narasimhan et al., 2017], and an exponentially-distributed recombination rate with mean 10^−8^ per site per generation, with maximum value truncated at 3 × 10^−8^, as in Schrider and Kern [2017] and Mughal and DeGiorgio [2019]. For *D. melanogaster*, our recombination rate was a uniform 5 × 10^−9^ per site per generation (equivalent to 5 × 10^−7^ cM per base), and our mutation rate was 10^−9^ per site per generation, as in Garud et al. [2015] and Harris et al. [2018b].

Our simulated human demographic histories consisted of European-descended CEU models and sub-Saharan African YRI models. The CEU model features a severe bottleneck reducing population effective size by an order of magnitude approximately 2000 generations before sampling, followed by a population expansion over two orders of magnitude leading to present day. The YRI model does not contain severe bottlenecks, and also includes an expansion similarly to the CEU model. Thus, the simulated CEU population has an approximately twofold reduction in its level of background genetic diversity relative to the YRI. For each of 1000 replicates under each power evaluation experiment (see below), we generated a simulated chromosome in SLiM of length 500 kilobases (kb) and scanned it with a sliding analysis window of size 117 SNPs, advancing by 12 SNPs per iteration. A window of 117 SNPs roughly corresponds to the number of SNPs expected in a physical window of size 40 kb for our sample size of 100 European diploid individuals, or 20 kb for 100 sub-Saharan African diploid individuals [Watterson, 1975]. We selected this window size because it is over this interval that pairwise linkage disequilibrium (LD) between SNPs decays by more than one-third on average in the human genome [Jakobsson et al., 2008]. This makes it unlikely that elevated values of the *T* statistic are due to background LD. For each analysis window of each neutral replicate, we generated a normalized, descending-order haplotype frequency spectrum truncated at a particular *K* between 10 and 25, and took the average of this truncated spectrum across all windows to produce an estimate of the neutral spectrum to create a baseline for variation in the absence of a selective sweep.

We simulated the *Drosophila* Genetic Reference Panel [DGRP; Mackay et al., 2012] *D. melanogaster* demographic history following the protocol of Harris et al. [2018b], adapting the model of Duchen et al. [2013] (Figure S11). Here, an ancestral African population (effective size *N*_1_) experiences a bottleneck at time *T*_B_, contracting to size *N*_B_ for 1000 generations before expanding to size 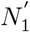. After the bottleneck, the ancestral European population diverges from the ancestral African population at time *τ*_1_, and begins with an effective size *N*_2_. The European population grows exponentially to its modern size, 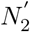. At time *τ*_2_, the North American population ancestral to the modern DGRP sample is generated with initial size from the admixture of the European and African populations, modeled as a single event, with a proportion *α* of North American genomes deriving from African ancestors, and a proportion (1 – *α*) deriving from European ancestors, where *α* < 1/2. The North American population grows exponentially to its final size, 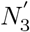. We draw each of the aforementioned model parameters according to their posterior probability density (Harris et al. [2018b], Table S1), thereby incorporating uncertainty into the demographic model. We used analysis windows of size 400 SNPs and a step size of 40 SNPs for *D. melanogaster* simulations. This represents the expected number of SNPs in a 10 kb window, over which a pairwise LD decay of greater than one-third occurs for the DGRP dataset [Garud et al., 2015].

To assess the power of the *T* statistic to differentiate between neutrality and sweeps, we simulated a variety of selective sweep scenarios, defined primarily by their combination of selection time *t*, selection strength *s*, and number of initially sweeping haplotypes *ν*. For human models, we simulated selection on *de novo* mutations arising at times *t* ∈ {200, 500,1000,1500, 2000, 4000} generations prior to sampling. We simulated sweeps on *ν* ∈ {1, 2, 4, 8,16, 32} distinct sweeping haplotypes, and chose selection coefficients uniformly at random following three schemes of weak, mixed, and strong selection coefficients. Weak selection was for *s* ∈ [0.005,0.05] and strong was for *s* ∈ [0.05,0.5], with mixed comprising both scenarios, *s* ∈ [0.005, 0.5] drawn uniformly at random specifically from a log-scale. We maintained window sizes identical to those for analysis of neutral replicates, this time generating a spectrum of normalized counts for each haplotype in each window. We computed the *T* statistic for each window and kept the largest for a replicate as its score. We also computed the score in this manner for each existing neutral replicate. We assessed the power of our approach for each parameter set as the proportion of sweep replicates whose score exceeded the top 1% or 5% of scores under neutrality.

To evaluate background selection as a potential confounding factor in identifying selective sweeps, we performed simulations in which we allowed for deleterious mutations to arise within the simulated chromosome throughout the simulation while maintaining all other parameters identical to neutrality. Our protocol was similar to that of Harris et al. [2018a], and covered the human CEU and YRI models. As with our previous simulations, we generated a genomic region of length 500 kb with identical mutation rate and population sizes as previously, evolving once again for a duration of 20N generations (*N* = 10^4^ diploids, the effective size during the burn-in period). At the center of the simulated sequence, we introduced a gene of length either 11 kb (small) or 55 kb (large) consisting of a 5’ untranslated region (UTR, length 200 bases), either 10 (small) or 50 (large) exons (100 bases each) and nine (small) or 49 (large) introns (one kb each) alternating for 10 (small) or 54 (large) kb, and a 3’ UTR (800 bases). We based the sizes of genetic elements on human genome-wide mean values [Mignone et al., 2002, Sakharkar et al., 2004]. Within the gene, strongly deleterious mutations (*s* = —0.1; gamma distribution of fitness effects with shape parameter 0.2) arose at rates of 50% within the UTRs, 75% within the exons, and 10% within the introns, while all other mutations within the gene and across the rest of the chromosome were selectively neutral. To enhance the effect of background selection under this scenario, we reduced the recombination rate from r = 10^−8^ to *r* = 10^−10^ per site per generation within the central gene.

Finally, we applied the *T* statistic to human empirical data from the 1000 Genomes Project [Auton et al., 2015], as well as to the DGRP inbred *D. melanogaster* dataset [Mackay et al., 2012]. The former application served primarily as a validation of our approach, as positive selection in the human genome has been widely explored. The latter application represented a typical insect model system that has also been well studied and diverges in size, genome architecture, and population history from humans. Our protocols for analyzing either dataset were identical in approach. For each, we applied a sliding window to all autosomes in the subject genome, basing window size on the interval over which LD, measured as *r*^2^, decayed below one-third of its original value relative to pairs of loci separated by one kb. This matched the prior approaches of [Garud et al., 2015, Harris et al., 2018a]. For humans, our window was size 117 SNPs, and for *D. melanogaster*, it was 400 SNPs, both matching our values for simulation experiments. Following the scans of CEU and YRI, we filtered windows overlapping chromosomal regions of low alignability and mappability, removing windows overlapping with chromosomal regions of mean CRG100 score less than 0.9. For *D. melanogaster*, we removed strains 49, 85, 101, 109, 136, 153, 237, 309, 317, 325, 338, 352, 377, 386, 426, 563, and 802 from our analysis due to their high number of heterozygous sites, and treated remaining heterozygous sites as missing data, as in Garud et al. [2015].

We then intersected the locations of computed *T* statistic values with the coordinates for protein- and RNA-coding genes based on hg19 and Dmel 5.13 annotations for humans and *D. melanogaster*, respectively. We assigned *p*-values to the 40 genes with the largest associated values of *T* by generating 10^6^ neutral replicates simulated in ms [Hudson, 2002] under demographic models inferred with `smc++` [Terhorst et al., 2017] for humans, and based on the Duchen et al. [2013] model for *D. melanogaster*, drawing parameters as previously. For each replicate, we simulated a sequence of length drawn uniformly at random from the set of all gene lengths, appended with the minimum number of nucleotides necessary to allow the application of full analysis windows centered across the entire length of the simulated gene. As an example, for a simulated human gene of length *L* nucleotides, we appended additional sequence length guaranteeing that 117-SNP windows centered at the first SNP and the last SNP of the simulated gene could be constructed. This allowed us to obtain a *T* statistic for at least one whole analysis window centered on the simulated gene during each replicate. The *p*-value for a selection candidate is the proportion of *T* statistics across all neutral replicates (using the maximum value for a replicate if there was more than one analysis window) that exceeded the maximum *T* associated with the candidate. All *p*-values were Bonferroni corrected for multiple testing [Neyman and Pearson, 1928], where a significant *p*-value was *p* < 0.05/*G* and where *G* is the number of genes for which we assigned a score in the organism. Accordingly, *G*_human_ = 18, 785 and phuman = 2.6617 × 10^−6^ for humans, whereas *G*_Dm_ = 10,000 and *p*_Dm_ = 5 × 10^−6^ for *D. melanogaster*.

## Acknowledgments

This work was funded by National Institutes of Health grant R35GM128590, by National Science Foundation grant DEB-1753489, and by the Alfred P. Sloan Foundation. Portions of this research were conducted with Advanced CyberInfrastructure computational resources provided by the Institute for CyberScience at Pennsylvania State University.

