## Supplementary material for "A likelihood approach for uncovering selective sweep signatures from haplotype data": Tables S1-S3 and Figures S1-S17

Table S1: Top 40 sweep candidates at RNA- and protein-coding genes of the Central European (CEU) human population, for both haplotype and multilocus genotype (MLG) data. Candidates presented are those that remained after filtering for mappability and alignability (see *Materials and Methods*), together with associated $T$ statistics and inferred number of sweeping haplotypes $\widehat{m}$. Target genes that pass the significance threshold are colored in gold in the "$p$-value" columns. Genes whose sweeps are assigned as hard ($\widehat{m} = 1$) are shaded in red in the "Inferred $\widehat{m}$" columns, while soft sweeps ($\widehat{m} \geq 2$) are colored in blue.

| | Top gene (hap) | Maximum $T$ (hap) | Inferred $\widehat{m}$ (hap) | $p$-value (hap) | Top gene (MLG) | Maximum $T$ (MLG) | Inferred $\widehat{m}$ (MLG) | $p$-value (MLG) |
|---|---|---|---|---|---|---|---|---|
| 1 | *ZRANB3* | 283.0868 | 1 | $< 10^{-6}$ | *LCT* | 144.63775 | 1 | $< 10^{-6}$ |
| 2 | *ZNF546* | 273.0766 | 1 | $1.0\times10^{-6}$ | *ZNF546* | 129.42416 | 1 | $1.0\times10^{-6}$ |
| 3 | *LCT* | 270.4884 | 1 | $2.0\times10^{-6}$ | *XIRP2* | 125.43142 | 2 | $4.0\times10^{-6}$ |
| 4 | *DARS* | 263.5470 | 1 | $2.0\times10^{-6}$ | *RSPH3* | 124.44902 | 1 | $4.0\times10^{-6}$ |
| 5 | *AC093391.2* | 260.7492 | 1 | $3.0\times10^{-6}$ | *MCM6* | 121.17195 | 1 | $1.0\times10^{-5}$ |
| 6 | *RSPH3* | 259.8491 | 1 | $3.0\times10^{-6}$ | *DARS* | 119.99953 | 1 | $1.1\times10^{-5}$ |
| 7 | *AC005592.1* | 257.5720 | 1 | $3.0\times10^{-6}$ | *HLA−DPB1* | 119.14111 | 1 | $1.1\times10^{-5}$ |
| 8 | *MCM6* | 254.2586 | 1 | $3.0\times10^{-6}$ | *AC005592.1* | 118.34704 | 1 | $1.1\times10^{-5}$ |
| 9 | *UBXN4* | 248.8716 | 1 | $7.0\times10^{-6}$ | *ZRANB3* | 117.16269 | 1 | $1.4\times10^{-5}$ |
| 10 | *BCAS3* | 241.7578 | 1 | $1.2\times10^{-5}$ | *SFPQ* | 113.90790 | 1 | $1.7\times10^{-5}$ |
| 11 | *SFPQ* | 241.4927 | 1 | $1.2\times10^{-5}$ | *BCAS3* | 113.88786 | 2 | $1.7\times10^{-5}$ |
| 12 | *HLA−DPB1* | 234.8464 | 1 | $1.7\times10^{-5}$ | *UBXN4* | 110.94659 | 1 | $2.0\times10^{-5}$ |
| 13 | *EP400NL* | 226.1789 | 1 | $3.0\times10^{-5}$ | *COL5A2* | 110.01764 | 2 | $2.2\times10^{-5}$ |
| 14 | *UNC5D* | 222.8515 | 1 | $3.7\times10^{-5}$ | *DGKI* | 110.00258 | 1 | $2.2\times10^{-5}$ |
| 15 | *DGKI* | 222.0198 | 1 | $3.8\times10^{-5}$ | *EP400NL* | 109.60544 | 1 | $2.4\times10^{-5}$ |
| 16 | *PSMB2* | 221.9329 | 1 | $3.9\times10^{-5}$ | *R3HDM1* | 103.65978 | 1 | $4.9\times10^{-5}$ |
| 17 | *CCDC178* | 218.1099 | 1 | $5.0\times10^{-5}$ | *DIP2C* | 100.41876 | 2 | $6.8\times10^{-5}$ |
| 18 | *C9orf3* | 216.5134 | 1 | $5.6\times10^{-5}$ | *EYS* | 98.21013 | 2 | $7.8\times10^{-5}$ |
| 19 | *R3HDM1* | 215.3070 | 1 | $5.7\times10^{-5}$ | *UNC5D* | 97.25789 | 1 | $8.3\times10^{-5}$ |
| 20 | *RAB3GAP1* | 212.4465 | 1 | $6.0\times10^{-5}$ | *STK32B* | 96.54516 | 2 | $8.8\times10^{-5}$ |
| 21 | *XIRP2* | 210.6209 | 1 | $6.9\times10^{-5}$ | *FGL1* | 95.42930 | 2 | $9.4\times10^{-5}$ |
| 22 | *COL5A2* | 208.3025 | 2 | $7.8\times10^{-5}$ | *BNC2* | 94.96477 | 1 | $9.7\times10^{-5}$ |
| 23 | *WWOX* | 207.7392 | 1 | $8.0\times10^{-5}$ | *AC093391.2* | 94.87813 | 1 | $9.7\times10^{-5}$ |
| 24 | *HLA−DRB5* | 198.7980 | 1 | $1.08\times10^{-4}$ | *SMCO2* | 93.85338 | 1 | $1.04\times10^{-4}$ |
| 25 | *ZNF211* | 198.0592 | 1 | $1.15\times10^{-4}$ | *HLA−DPA1* | 92.28441 | 2 | $1.16\times10^{-4}$ |
| 26 | *EYS* | 197.2657 | 1 | $1.17\times10^{-4}$ | *KCNQ5* | 92.27282 | 2 | $1.16\times10^{-4}$ |
| 27 | *SYNE2* | 195.3472 | 1 | $1.32\times10^{-4}$ | *SYNE2* | 92.13772 | 1 | $1.19\times10^{-4}$ |
| 28 | *MIR548AZ* | 195.3472 | 1 | $1.32\times10^{-4}$ | *MIR548AZ* | 92.13772 | 1 | $1.19\times10^{-4}$ |
| 29 | *KCNQ5* | 195.2536 | 1 | $1.33\times10^{-4}$ | *SLC12A1* | 91.19790 | 1 | $1.26\times10^{-4}$ |
| 30 | *PPARD* | 191.6392 | 1 | $1.51\times10^{-4}$ | *DNAH14* | 89.58234 | 1 | $1.42\times10^{-4}$ |
| 31 | *ALG12* | 191.5975 | 1 | $1.51\times10^{-4}$ | *TMEM232* | 86.93562 | 1 | $1.75\times10^{-4}$ |
| 32 | *ACER1* | 191.3206 | 1 | $1.52\times10^{-4}$ | *SCP2* | 86.33526 | 1 | $1.81\times10^{-4}$ |
| 33 | *SCP2* | 189.0051 | 1 | $1.65\times10^{-4}$ | *SETX* | 86.01296 | 1 | $1.88\times10^{-4}$ |
| 34 | *SPATA6L* | 188.8695 | 1 | $1.66\times10^{-4}$ | *POLN* | 85.87041 | 1 | $1.89\times10^{-4}$ |
| 35 | *SMCO2* | 187.0748 | 1 | $1.76\times10^{-4}$ | *MYO18B* | 85.33800 | 2 | $1.93\times10^{-4}$ |
| 36 | *ZNF780B* | 186.6733 | 1 | $1.76\times10^{-4}$ | *CPPED1* | 85.26133 | 1 | $1.94\times10^{-4}$ |
| 37 | *GAB2* | 186.1554 | 1 | $1.79\times10^{-4}$ | *PSMB2* | 84.80460 | 1 | $2.04\times10^{-4}$ |
| 38 | *ECD* | 185.8040 | 1 | $1.82\times10^{-4}$ | *MYO9A* | 84.15219 | 1 | $2.18\times10^{-4}$ |
| 39 | *MYO9A* | 185.3759 | 1 | $1.88\times10^{-4}$ | *RAB3GAP1* | 82.50744 | 1 | $2.49\times10^{-4}$ |
| 40 | *NCKAP5L* | 182.1102 | 2 | $2.11\times10^{-4}$ | *ZNF780B* | 81.97445 | 1 | $2.56\times10^{-4}$ |

Table S2: Top 40 sweep candidates at RNA- and protein-coding genes of the sub-Saharan Yoruban (YRI) human population, for both haplotype and multilocus genotype (MLG) data. Candidates presented are those that remained after filtering for mappability and alignability (see *Materials and Methods*), together with associated $T$ statistics and inferred number of sweeping haplotypes $\widehat{m}$. Target genes that pass the significance threshold are colored in gold in the "$p$-value" columns. Genes whose sweeps are assigned as hard ($\widehat{m} = 1$) are shaded in red in the "Inferred $\widehat{m}$" columns, while soft sweeps ($\widehat{m} \geq 2$) are colored in blue.

| | Top gene (hap) | Maximum $T$ (hap) | Inferred $\widehat{m}$ (hap) | $p$-value (hap) | Top gene (MLG) | Maximum $T$ (MLG) | Inferred $\widehat{m}$ (MLG) | $p$-value (MLG) |
|---|---|---|---|---|---|---|---|---|
| 1 | *SPRED3* | 284.7154 | 1 | $< 10^{-6}$ | *ITGAE* | 130.88759 | 2 | $< 10^{-6}$ |
| 2 | *SYT1* | 275.5915 | 2 | $1.0\times10^{-6}$ | *SPRED3* | 124.28967 | 1 | $< 10^{-6}$ |
| 3 | *HLA−DPB2* | 249.4059 | 1 | $2.0\times10^{-6}$ | *SYT1* | 113.15402 | 2 | $< 10^{-6}$ |
| 4 | *ITGAE* | 243.4470 | 1 | $2.0\times10^{-6}$ | *HLA−DPB2* | 96.96212 | 3 | $4.0\times10^{-6}$ |
| 5 | *TLR5* | 239.7464 | 1 | $2.0\times10^{-6}$ | *TLR5* | 93.10259 | 1 | $5.0\times10^{-6}$ |
| 6 | *SUGCT* | 216.5221 | 1 | $5.0\times10^{-6}$ | *SUGCT* | 89.76026 | 2 | $5.0\times10^{-6}$ |
| 7 | *FAM60A* | 214.2499 | 1 | $5.0\times10^{-6}$ | *CTDSPL2* | 85.71732 | 2 | $5.0\times10^{-6}$ |
| 8 | *GTSF1* | 214.0371 | 1 | $5.0\times10^{-6}$ | *NNT* | 83.82059 | 3 | $7.0\times10^{-6}$ |
| 9 | *MIR548H3* | 211.5784 | 1 | $6.0\times10^{-6}$ | *FAM60A* | 83.81482 | 1 | $7.0\times10^{-6}$ |
| 10 | *ZFPM1* | 209.9309 | 1 | $7.0\times10^{-6}$ | *FAM98C* | 82.72407 | 2 | $7.0\times10^{-6}$ |
| 11 | *NNT* | 206.8125 | 1 | $7.0\times10^{-6}$ | *CSMD3* | 80.82570 | 3 | $1.0\times10^{-5}$ |
| 12 | *NANS* | 206.6424 | 2 | $7.0\times10^{-6}$ | *PSCA* | 80.48072 | 3 | $1.0\times10^{-5}$ |
| 13 | *MGAT4A* | 198.0702 | 1 | $1.1\times10^{-5}$ | *MITF* | 79.66823 | 2 | $1.1\times10^{-5}$ |
| 14 | *SEMA3C* | 195.9268 | 1 | $1.2\times10^{-5}$ | *CNGA3* | 79.64351 | 1 | $1.1\times10^{-5}$ |
| 15 | *CNGA3* | 195.1357 | 1 | $1.2\times10^{-5}$ | *PAWR* | 77.90110 | 1 | $2.0\times10^{-5}$ |
| 16 | *MIR548AE2* | 192.3556 | 1 | $1.4\times10^{-5}$ | *PTPRT* | 77.61271 | 1 | $2.0\times10^{-5}$ |
| 17 | *LONP2* | 192.3556 | 1 | $1.4\times10^{-5}$ | *MIR548H3* | 76.77193 | 1 | $2.0\times10^{-5}$ |
| 18 | *HEMGN* | 190.1230 | 1 | $1.5\times10^{-5}$ | *NRXN3* | 76.63015 | 1 | $2.0\times10^{-5}$ |
| 19 | *GBA3* | 190.0849 | 1 | $1.5\times10^{-5}$ | *CCBL1* | 74.59808 | 2 | $2.4\times10^{-5}$ |
| 20 | *SDS* | 187.9896 | 2 | $1.6\times10^{-5}$ | *GTSF1* | 74.39620 | 3 | $2.4\times10^{-5}$ |
| 21 | *HIF1AN* | 185.7172 | 1 | $1.7\times10^{-5}$ | *DKK2* | 74.27090 | 4 | $2.4\times10^{-5}$ |
| 22 | *PREX1* | 185.2450 | 1 | $1.7\times10^{-5}$ | *RBBP4* | 72.89650 | 3 | $2.6\times10^{-5}$ |
| 23 | *COX7B2* | 184.4306 | 3 | $1.7\times10^{-5}$ | *ABCA17P* | 72.48000 | 2 | $2.9\times10^{-5}$ |
| 24 | *ABCA17P* | 182.5478 | 2 | $1.9\times10^{-5}$ | *SHPK* | 72.33416 | 4 | $3.0\times10^{-5}$ |
| 25 | *RGS18* | 182.3130 | 2 | $2.0\times10^{-5}$ | *HIF1AN* | 72.16977 | 3 | $3.0\times10^{-5}$ |
| 26 | *LINC00506* | 181.7588 | 2 | $2.0\times10^{-5}$ | *HLA−DRB5* | 72.05417 | 1 | $3.0\times10^{-5}$ |
| 27 | *FAM98C* | 179.6052 | 2 | $2.5\times10^{-5}$ | *CNNM2* | 71.89649 | 3 | $3.0\times10^{-5}$ |
| 28 | *PTPRT* | 176.1634 | 1 | $2.5\times10^{-5}$ | *PPM1L* | 71.07533 | 1 | $3.2\times10^{-5}$ |
| 29 | *DKK2* | 174.5704 | 2 | $2.7\times10^{-5}$ | *CATSPERG* | 70.70611 | 4 | $3.6\times10^{-5}$ |
| 30 | *ADCY1* | 174.5669 | 1 | $2.7\times10^{-5}$ | *HLA−DPA1* | 70.68163 | 3 | $3.6\times10^{-5}$ |
| 31 | *CCBL1* | 174.1857 | 2 | $2.7\times10^{-5}$ | *CEACAM18* | 70.39460 | 4 | $3.8\times10^{-5}$ |
| 32 | *F11R* | 173.8417 | 1 | $2.7\times10^{-5}$ | *PTCHD4* | 70.10061 | 1 | $3.8\times10^{-5}$ |
| 33 | *HLA−DRB5* | 171.7244 | 1 | $3.1\times10^{-5}$ | *LMO7* | 70.07955 | 4 | $3.8\times10^{-5}$ |
| 34 | *RAB28* | 171.3009 | 1 | $3.1\times10^{-5}$ | *HECW1* | 69.96530 | 1 | $3.9\times10^{-5}$ |
| 35 | *BTNL2* | 167.7484 | 6 | $3.6\times10^{-5}$ | *AC011738.4* | 69.83389 | 4 | $3.9\times10^{-5}$ |
| 36 | *CSMD3* | 166.5819 | 1 | $3.6\times10^{-5}$ | *CASC4* | 69.02636 | 2 | $4.3\times10^{-5}$ |
| 37 | *CNNM2* | 165.5460 | 3 | $3.8\times10^{-5}$ | *ADCY1* | 68.73247 | 1 | $4.3\times10^{-5}$ |
| 38 | *CELF5* | 165.3101 | 1 | $4.0\times10^{-5}$ | *MIR548AE2* | 68.68579 | 1 | $4.4\times10^{-5}$ |
| 39 | *CNTNAP2* | 165.1435 | 1 | $4.1\times10^{-5}$ | *LONP2* | 68.68579 | 1 | $4.4\times10^{-5}$ |
| 40 | *INPP5A* | 163.5742 | 3 | $4.5\times10^{-5}$ | *COX7B2* | 68.67966 | 4 | $4.4\times10^{-5}$ |

Table S3: Top 40 sweep candidates at RNA- and protein-coding genes of the inbred North American DGRP *Drosophila melanogaster* population. Candidates presented are those that remained after filtering out individuals with excessive heterozygous sites (see *Materials and Methods*), together with associated $T$ statistics and inferred number of sweeping haplotypes $\widehat{m}$. Although no genes pass the significance threshold under the Duchen et al. [2013] model with parameter uncertainty, we include a "$p$-value" column for context. Genes whose sweeps are assigned as hard ($\widehat{m} = 1$) are shaded in red in the "Inferred $\widehat{m}$" columns, whereas soft sweeps ($\widehat{m} \geq 2$) are colored in blue.

| | Top gene | Maximum $T$ | Inferred $\widehat{m}$ | $p$–value |
|---|---|---|---|---|
| *1* | *CG11902* | 246.58748 | 1 | 0.003975 |
| *2* | *Ace* | 179.36445 | 1 | 0.021491 |
| *3* | *Uhg1* | 178.88792 | 1 | 0.021731 |
| *4* | *CG30047* | 163.07482 | 1 | 0.031653 |
| *5* | *CG8449* | 160.24925 | 1 | 0.033816 |
| *6* | *CG32473* | 156.88243 | 1 | 0.036437 |
| *7* | *Pimet* | 153.94931 | 3 | 0.039104 |
| *8* | *CG8774* | 152.28501 | 1 | 0.040431 |
| *9* | *CG8878* | 152.18730 | 3 | 0.040526 |
| *10* | *Ho* | 150.96343 | 1 | 0.041618 |
| *11* | *CG10669* | 148.29402 | 2 | 0.044246 |
| *12* | *CG6834* | 144.24962 | 2 | 0.048248 |
| *13* | *CG6830* | 143.68106 | 2 | 0.048856 |
| *14* | *ana3* | 142.79537 | 1 | 0.049806 |
| *15* | *CG11686* | 142.70239 | 1 | 0.049927 |
| *16* | *CG8378* | 136.96796 | 3 | 0.056287 |
| *17* | *rha* | 133.67244 | 1 | 0.060691 |
| *18* | *Su(var)3−7* | 129.54795 | 1 | 0.066547 |
| *19* | *Osi22* | 127.72869 | 2 | 0.069230 |
| *20* | *Rep3* | 125.14389 | 2 | 0.073284 |
| *21* | *wb* | 116.61486 | 1 | 0.087331 |
| *22* | *CG14715* | 115.31162 | 1 | 0.089483 |
| *23* | *CG17739* | 113.57495 | 3 | 0.092448 |
| *24* | *prp8* | 108.62382 | 3 | 0.101281 |
| *25* | *CG30049* | 107.44626 | 1 | 0.103421 |
| *26* | *CG6908* | 107.42041 | 1 | 0.103465 |
| *27* | *CG18476* | 107.30661 | 1 | 0.103703 |
| *28* | *Taf12* | 107.30661 | 1 | 0.103703 |
| *29* | *CG13177* | 107.27694 | 3 | 0.103766 |
| *30* | *Ravus* | 106.14619 | 1 | 0.105899 |
| *31* | *nompC* | 105.43706 | 2 | 0.107277 |
| *32* | *CG8773* | 104.05140 | 3 | 0.109950 |
| *33* | *CG12594* | 102.88848 | 1 | 0.112186 |
| *34* | *CG9510* | 102.39815 | 1 | 0.113153 |
| *35* | *CG9515* | 102.39815 | 1 | 0.113153 |
| *36* | *CG8508* | 101.20347 | 2 | 0.115509 |
| *37* | *Rab11* | 100.30730 | 1 | 0.117314 |
| *38* | *jar* | 99.29919 | 1 | 0.119355 |
| *39* | *CG8407* | 98.72060 | 3 | 0.120530 |
| *40* | *timeout* | 92.99725 | 1 | 0.133005 |

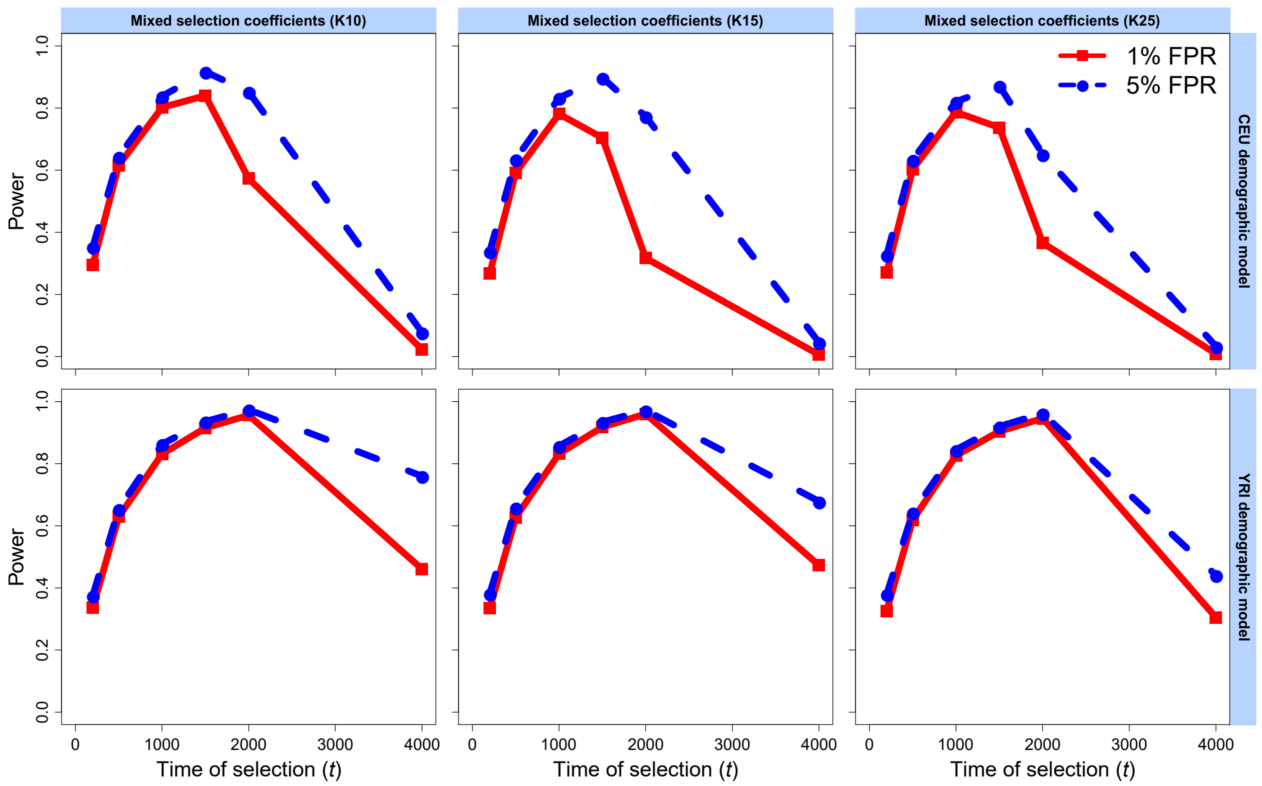


Figure S1: Power of the $T$ statistic at 1% and 5% false positive rates (FPRs) to detect hard selective sweeps from a single *de novo* mutation arising at time $t \in \{200, 500, 1000, 1500, 2000, 4000\}$ generations before sampling under the European CEU (top) and sub-Saharan African YRI (bottom) human demographic models. Mixed selection coefficients were drawn uniformly at random on a log-scale from $s \in [0.005, 0.5]$. Simulated replicates are identical to those in Figure 2, but with sample spectra of $K = 10$ (left), $K = 15$ (center), and $K = 25$ (right) most frequent haplotypes used for inference.

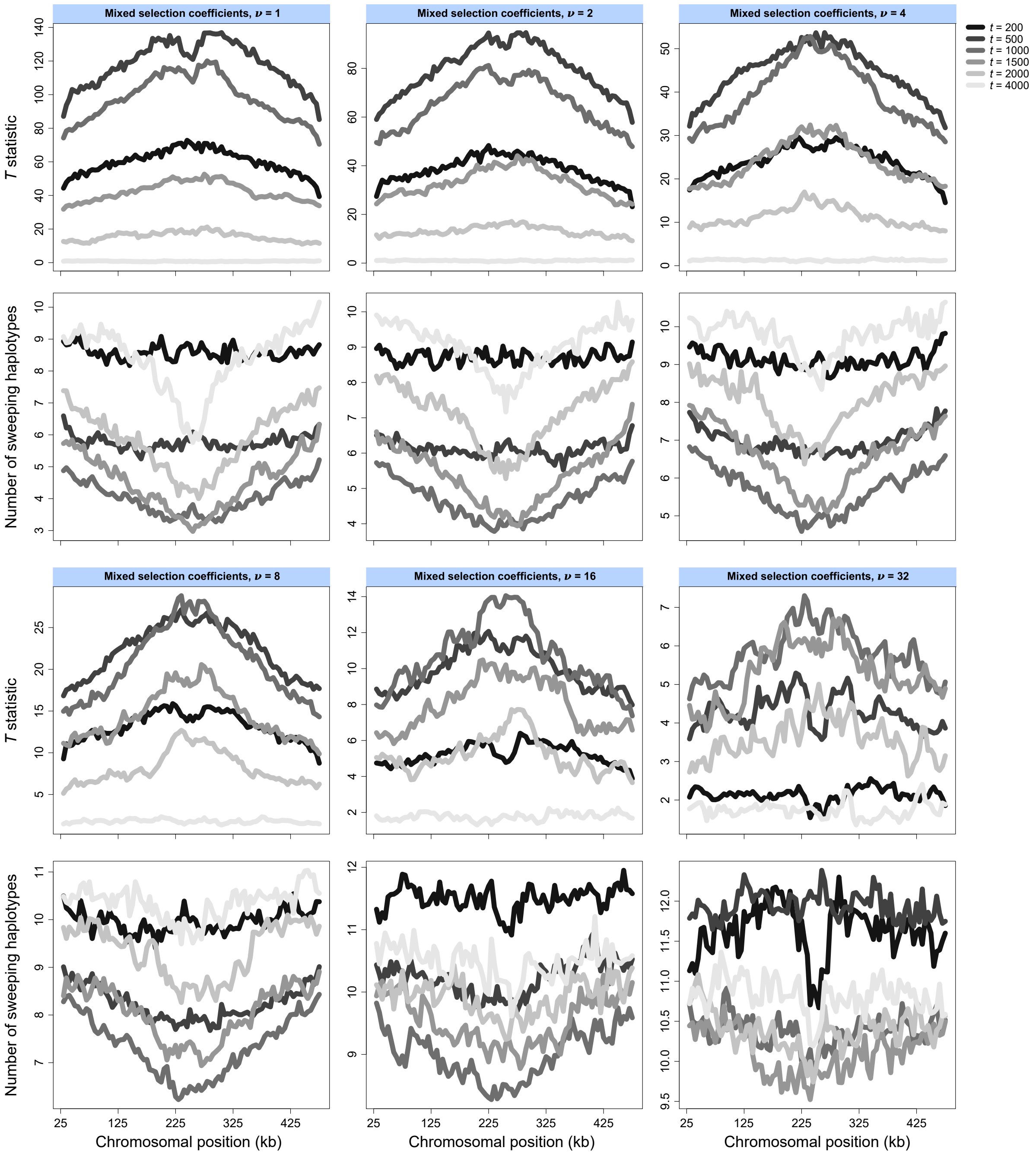


Figure S2: Mean spatial distribution of the $T$ statistic (first and third rows) and the inferred number of sweeping haplotypes ($\widehat{m}$; second and fourth rows) across the central 450 kb of a 500 kb chromosome simulated under the European CEU human demographic model. Each line is the average of 1000 simulated replicates initiated under identical selection parameters, consisting of mixed selection coefficients with $s \in [0.005, 0.5]$ drawn uniformly at random on a log-scale, times of selection $t \in \{200, 500, 1000, 1500, 2000, 4000\}$ generations prior to sampling, and number of sweeping haplotypes $\nu \in \{1, 2, 4, 8, 16, 32\}$. The simulated replicates here are identical to those in the top rows of Figures 2 and 3.

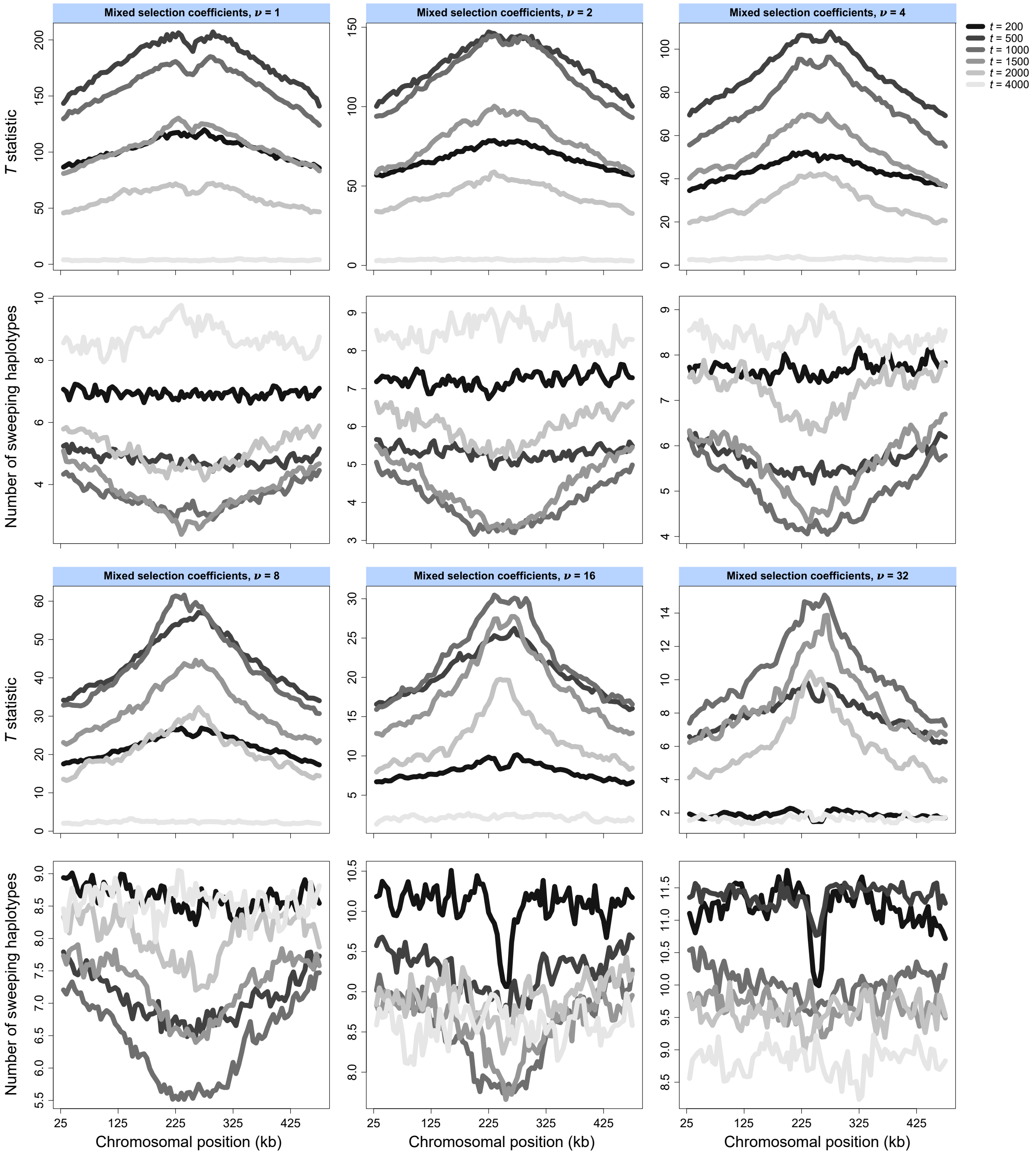


Figure S3: Mean spatial distribution of the $T$ statistic (first and third rows) and the inferred number of sweeping haplotypes ($\widehat{m}$; second and fourth rows) across the central 450 kb of a 500 kb chromosome simulated under the sub-Saharan African YRI human demographic model. Each line is the average of 1000 simulated replicates initiated under identical selection parameters, consisting of mixed selection coefficients with $s \in [0.005, 0.5]$ drawn uniformly at random on a log-scale, times of selection $t \in \{200, 500, 1000, 1500, 2000, 4000\}$ generations prior to sampling, and number of sweeping haplotypes $\nu \in \{1, 2, 4, 8, 16, 32\}$. The simulated replicates here are identical to those in the bottom rows of Figures 2 and 3.

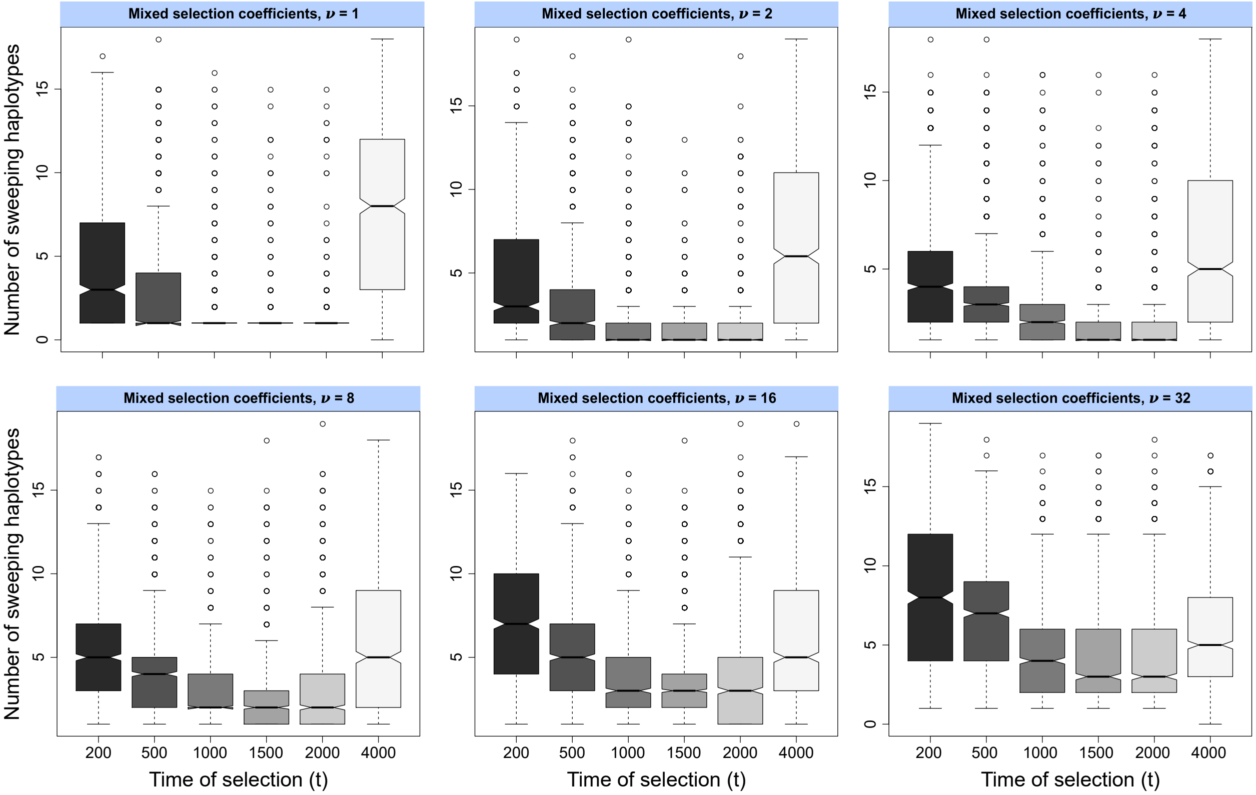


Figure S4: Box plots summarizing the distributions of the inferred number of sweeping haplotypes $\widehat{m}$ under the European CEU human demographic model for simulated selective sweeps with strength $s \in [0.005, 0.5]$ drawn uniformly at random on a log-scale and selection on $\nu \in \{1, 2, 4, 8, 16, 32\}$ distinct sweeping haplotypes. The $T$ statistic was computed from the $K = 20$ most frequent sampled haplotypes.

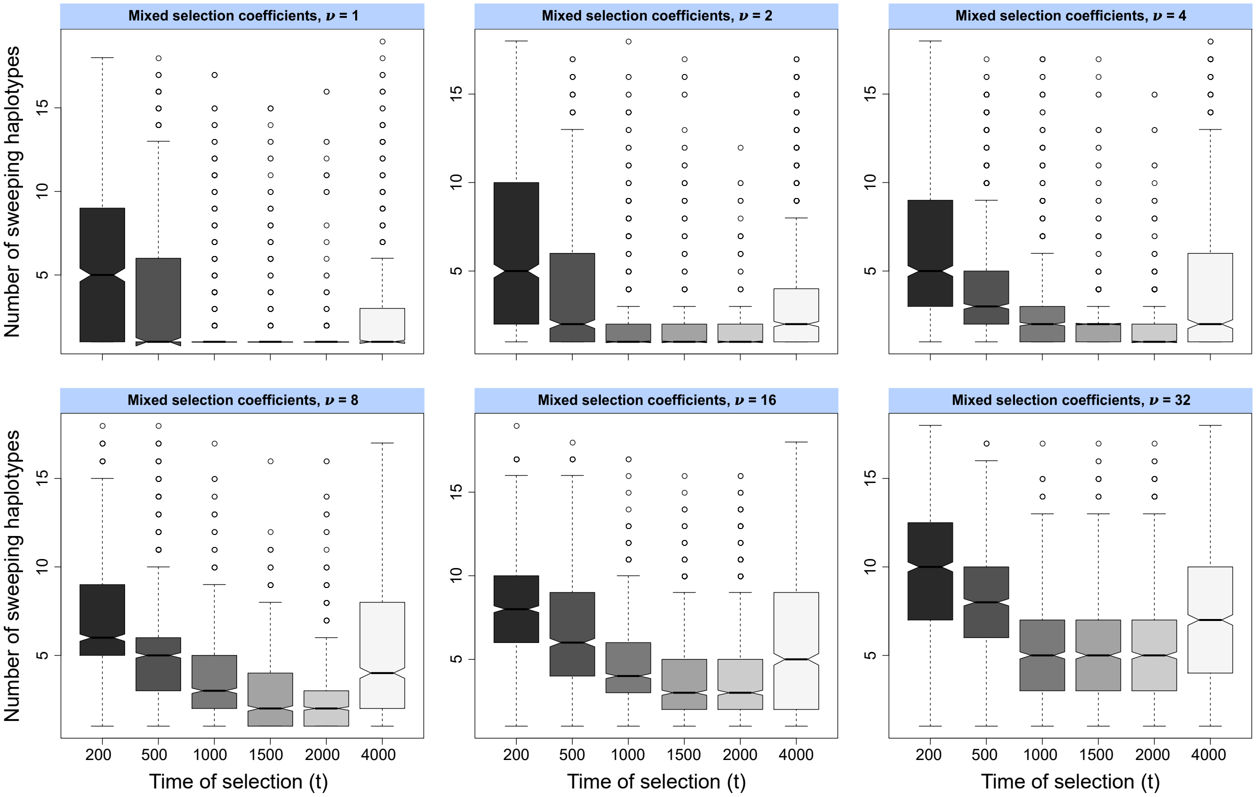


Figure S5: Box plots summarizing the distributions of the inferred number of sweeping haplotypes $\widehat{m}$ under the sub-Saharan African YRI human demographic model for simulated selective sweeps with strength $s \in [0.005, 0.5]$ drawn uniformly at random on a log-scale and selection on $\nu \in \{1, 2, 4, 8, 16, 32\}$ distinct sweeping haplotypes. The $T$ statistic was computed from the $K = 20$ most frequent sampled haplotypes.

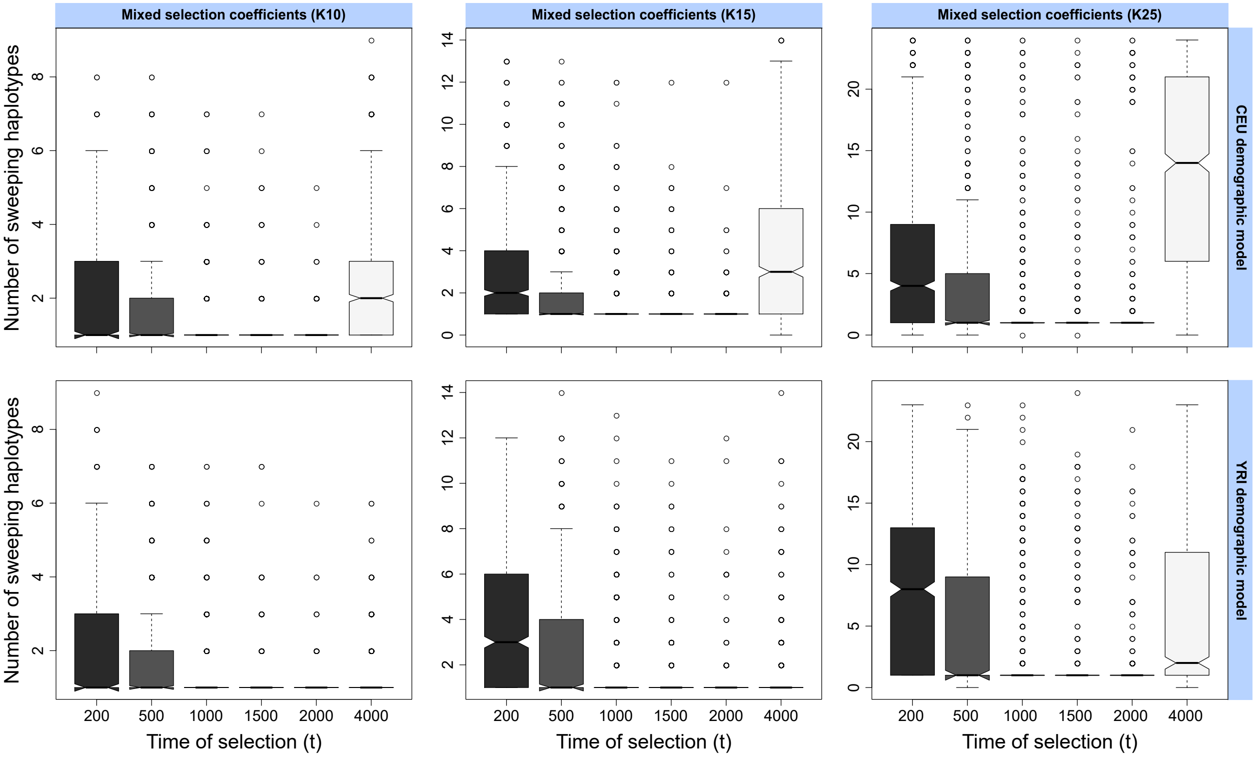


Figure S6: Box plots summarizing the distributions of the inferred number of sweeping haplotypes $\widehat{m}$ under the European CEU (top) and sub-Saharan African YRI (bottom) human demographic models for simulated hard selective sweeps with strength $s \in [0.005, 0.5]$ drawn uniformly at random on a log-scale. The $T$ statistic was computed from the $K = 10$ (left), $K = 15$ (center), or $K = 25$ (right) most frequent sampled haplotypes.

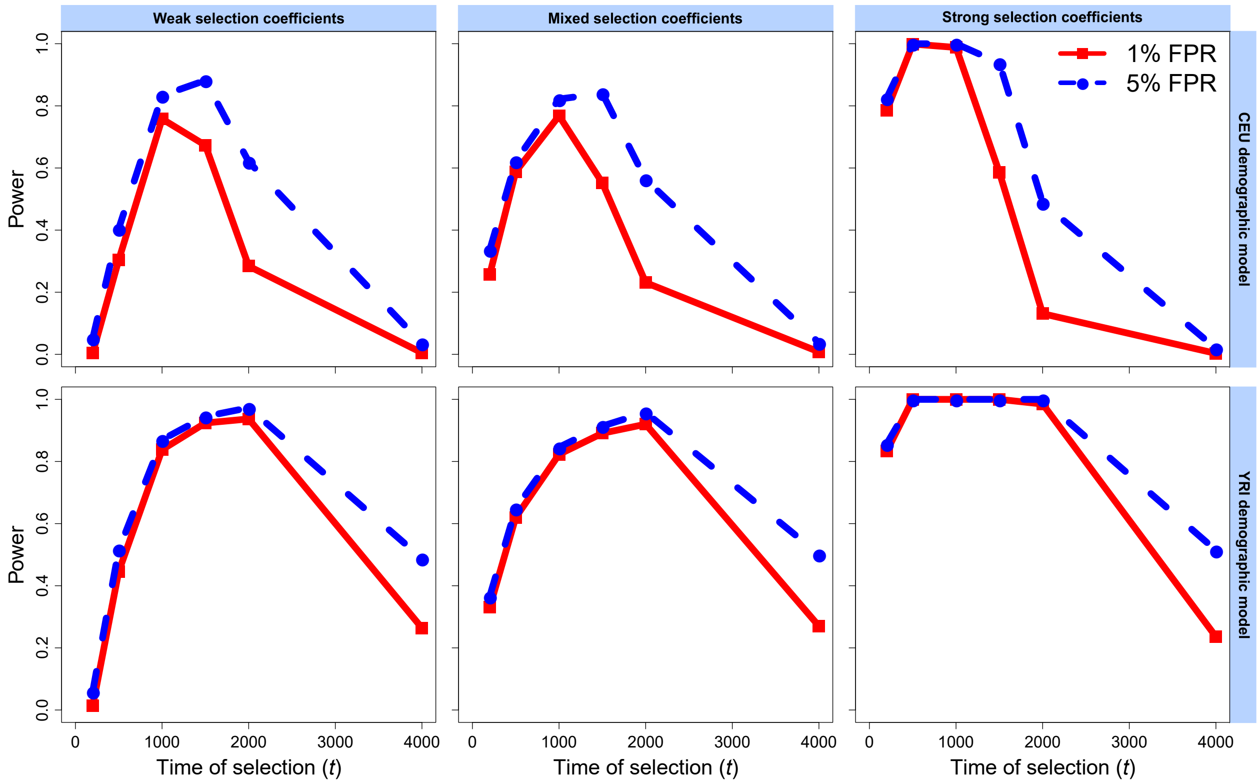


Figure S7: Power of the $T$ statistic at 1% and 5% false positive rates (FPRs) to detect hard selective sweeps from a single *de novo* mutation arising at time $t \in \{200, 500, 1000, 1500, 2000, 4000\}$ generations before sampling under the European CEU (top) and sub-Saharan African YRI (bottom) human demographic models, for unphased multilocus genotypes (MLGs). Simulated replicates are identical to those in Figures 2 and 3. Weak selection coefficients were drawn uniformly at random from $s \in [0.005, 0.05]$, strong selection coefficients were drawn uniformly at random from $s \in [0.05, 0.5]$, and mixed selection coefficients were drawn uniformly at random on a log-scale from $s \in [0.005, 0.5]$. All inferences used a spectrum of $K = 20$ for likelihood computations.

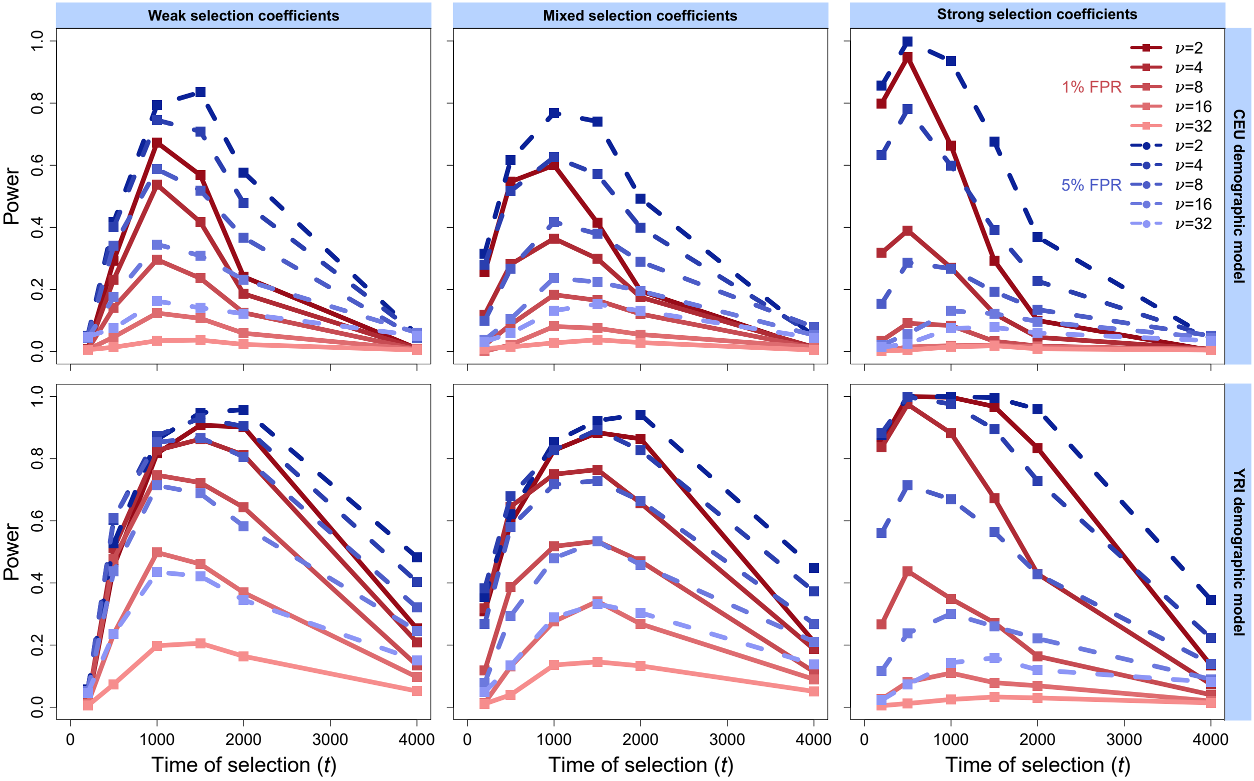


Figure S8: Power of the $T$ statistic at 1% and 5% false positive rates (FPRs) to detect soft selective sweeps from selection on standing variation on $\nu \in \{2, 4, 8, 16, 32\}$ distinct sweeping haplotypes beginning at time $t \in \{200, 500, 1000, 1500, 2000, 4000\}$ generations before sampling under the European CEU (top) and sub-Saharan African YRI (bottom) human demographic models, for unphased multilocus genotypes (MLGs). Simulated replicates are identical to those in Figures 2 and 3. Weak selection coefficients were drawn uniformly at random from $s \in [0.005, 0.05]$, strong selection coefficients were drawn uniformly at random from $s \in [0.05, 0.5]$, and mixed selection coefficients were drawn uniformly at random on a log-scale from $s \in [0.005, 0.5]$. All inferences used a spectrum of $K = 20$ for likelihood computations.

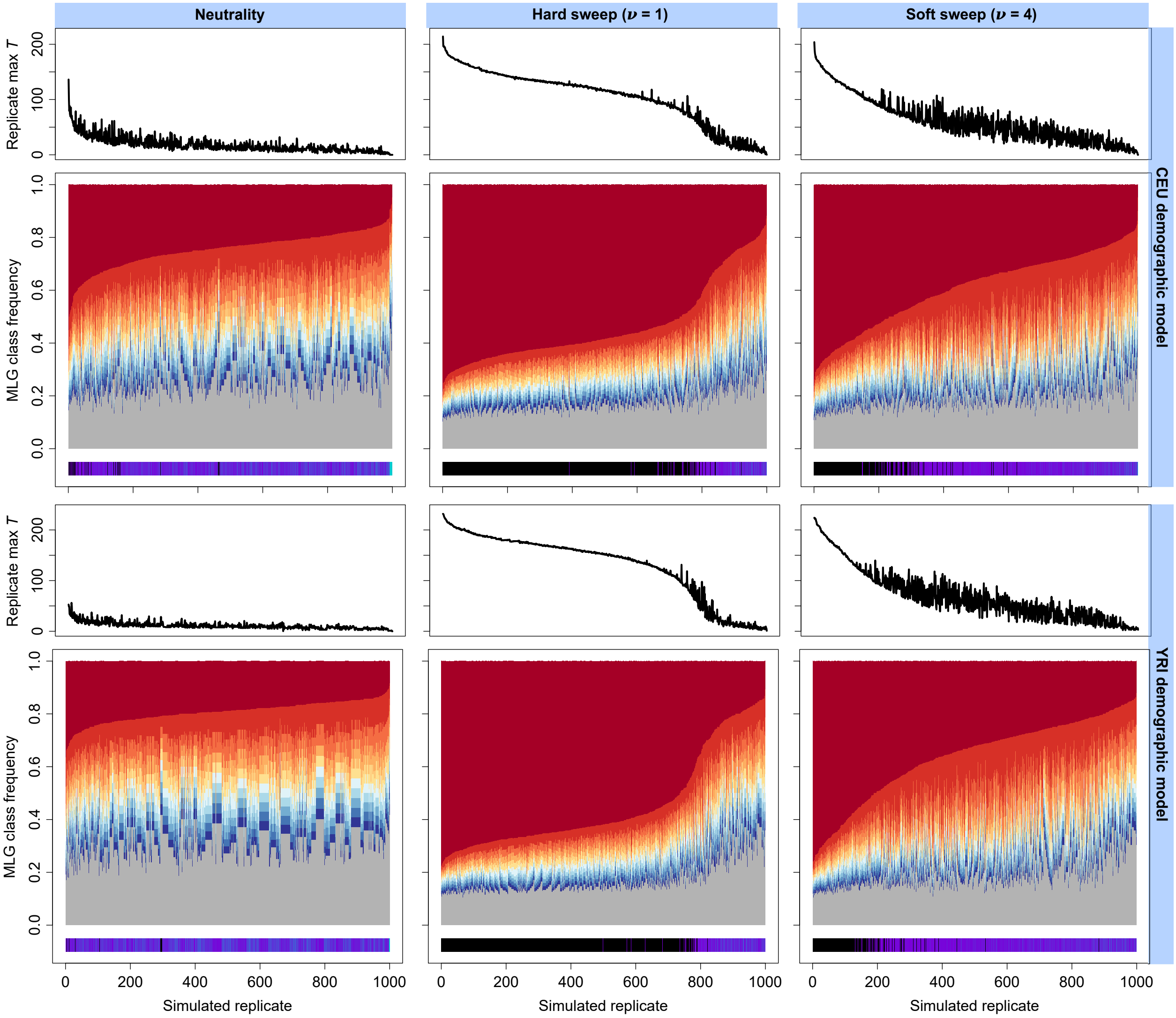


Figure S9: Truncated MLG frequency spectra ($K = 20$) across $10^3$ simulated replicates for analysis window of maximum replicate-wide $T$ statistic under neutral (left), hard sweep (center), and soft sweep (right) scenarios, for European CEU (top) and sub-Saharan African YRI (bottom) human demographic models. Each simulated replicate is one vertical slice within the greater plot, and the 10 most frequent MLGs are colored on a scale from red (most-frequent) to blue (10th most-frequent), while the remaining MLGs are shaded together in gray. Replicates are associated with their $T$ statistic (above) and their inferred $\widehat{m}$ (below). Inferred hard sweeps ($\widehat{m} = 1$) are indicated in black, whereas inferred soft sweeps ($\widehat{m} \geq 2$) are indicated on a color scale spanning purple (fewer sweeping MLGs) to teal (maximum of 20 sweeping MLGs, consistent with neutrality). Replicate spectra are arranged in decreasing order of most-frequent MLG frequency.

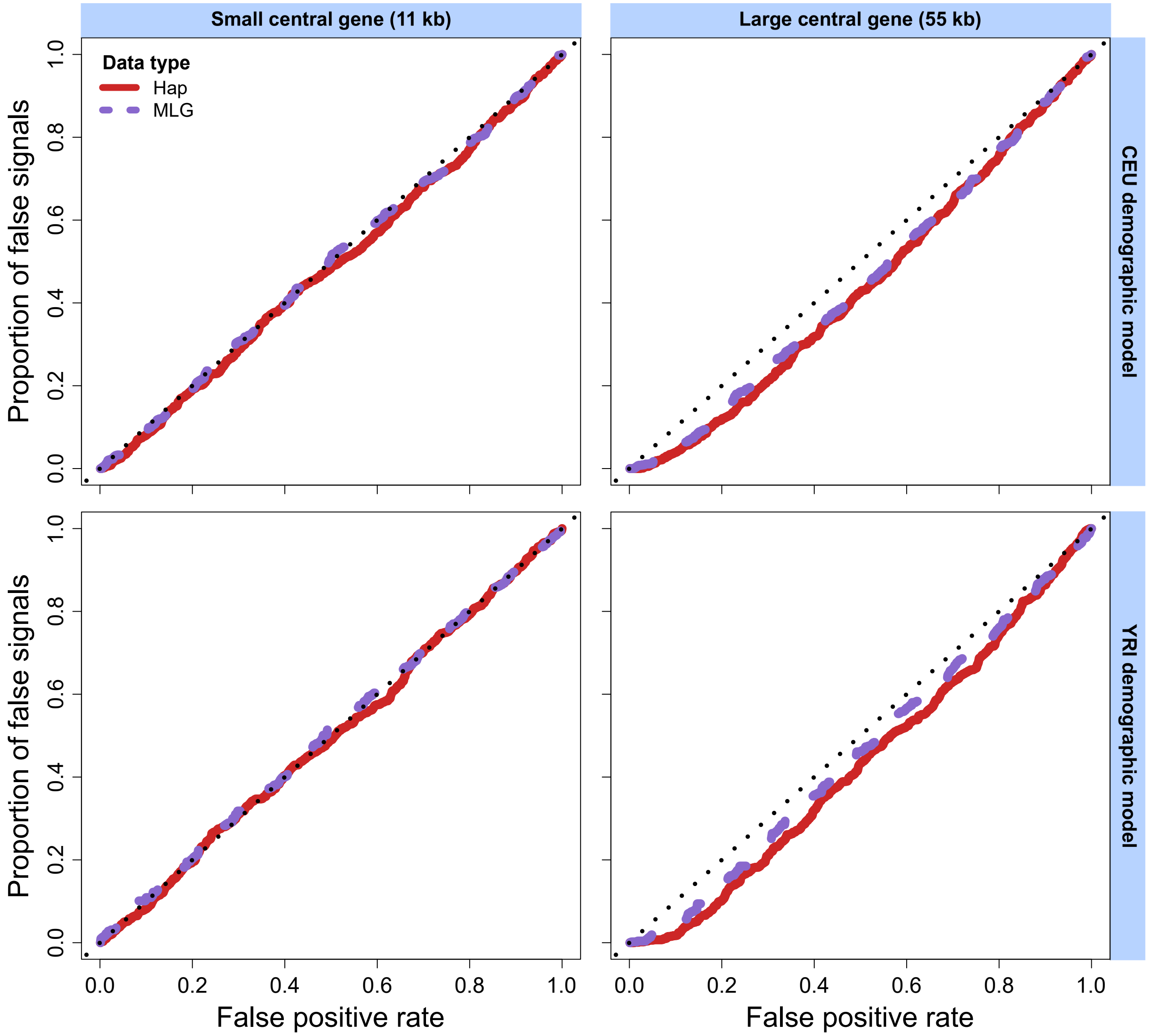


Figure S10: Effect of background selection on the distribution of the $T$ statistic relative to neutrality measured as the proportion of false signals under background selection as a function of the false positive rate under neutrality. Models considered are those for the human CEU (top) and YRI (bottom) populations, for background selection occurring on a central gene within a 500 kb simulated chromosome. Scenarios of a small 11 kb (left) or large 55 kb (right) central gene are considered across haplotype (hap, red) and multilocus genotype (MLG, purple) data.

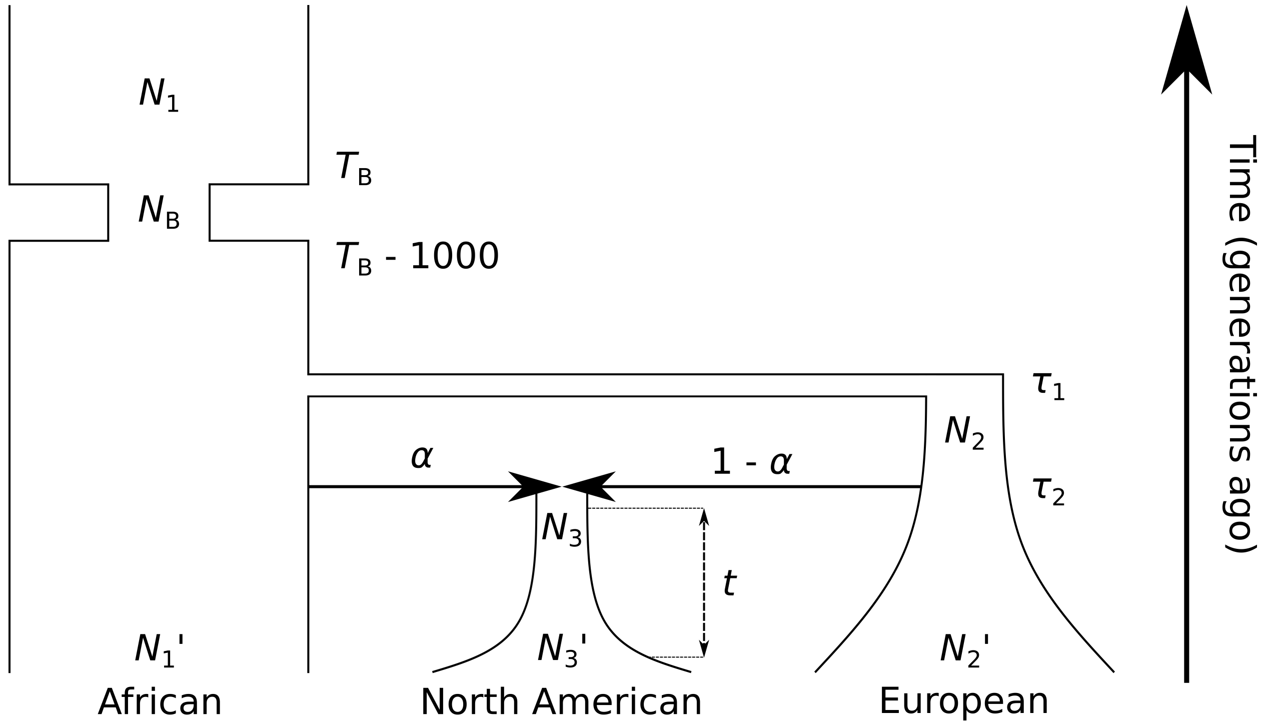


Figure S11: *D. melanogaster* demographic history model adapted from Duchen et al. [2013]. In this model, the modern DGRP [Mackay et al., 2012] North American *D. melanogaster* population descends from a recent admixture event between African and European ancestral populations. We used this model as the basis for all *D. melanogaster* simulations, drawing each parameter of the model from a posterior distribution, with probabilities as indicated in Table S1 of Harris et al. [2018b]. Because the order of events in this demographic history is fixed, we constrained that, starting from the present, we have $0 < t < \tau_2 < \tau_1 < T_{\mathrm{B}} - 1000 < T_{\mathrm{B}}$.

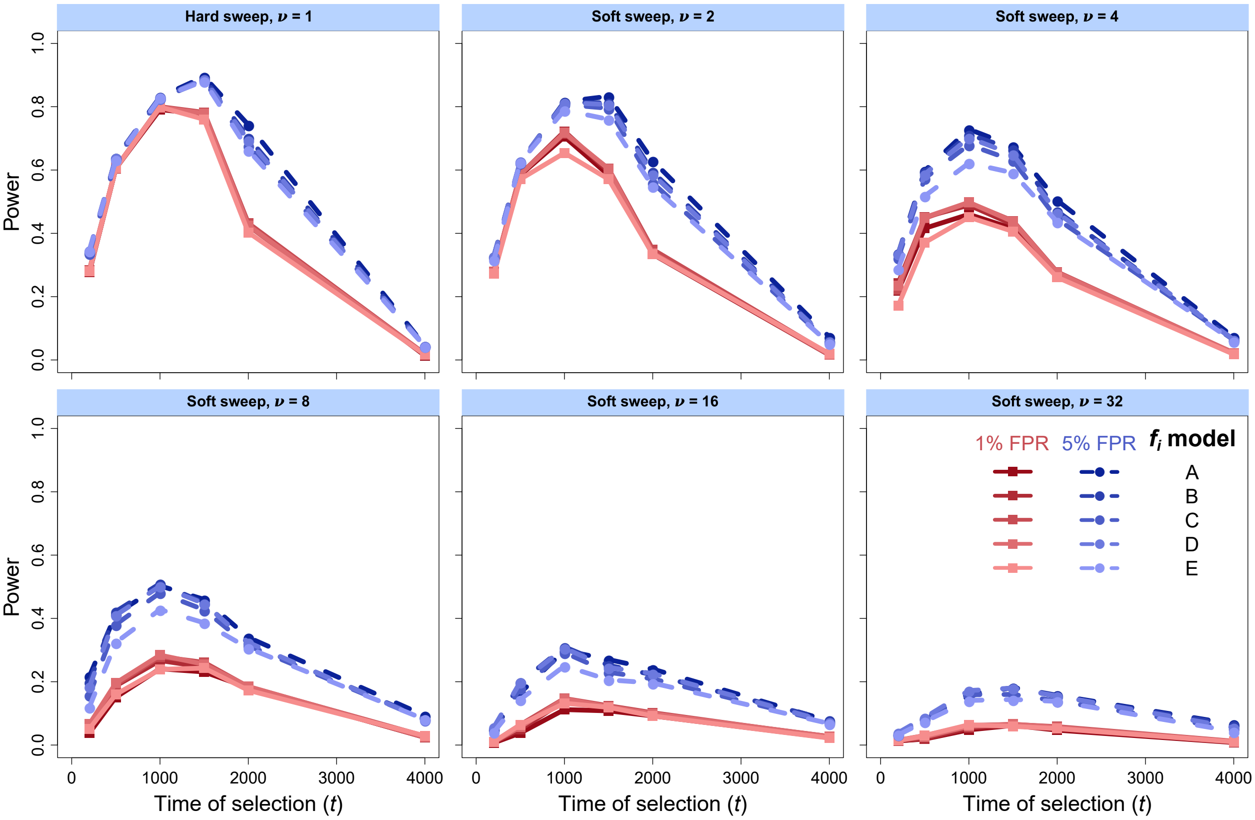


Figure S12: Powers of the $T$ statistic variants for different choices of $f_i$ at 1% and 5% false positive rates (FPRs) to detect selective sweeps on a CEU demographic history for haplotype frequency spectra truncated at $K = 20$ haplotypes. Analyzed data are identical to the CEU data represented in Figures 2 and 3. Models tested include uniform $f_i = 1/m$ (model A), $f_i = (1/i)/\sum_{j=1}^{m} 1/j$ (model B), $f_i = (1/i^2)/\sum_{j=1}^{m} 1/j^2$ (model C), $f_i = e^{-i}/\sum_{j=1}^{m} e^{-j}$ (model D), and $f_i = e^{-i^2}/\sum_{j=1}^{m} e^{-j^2}$ (model E). We chose $U = p_K$ and optimized $\varepsilon \in [0.0005, p_K]$.

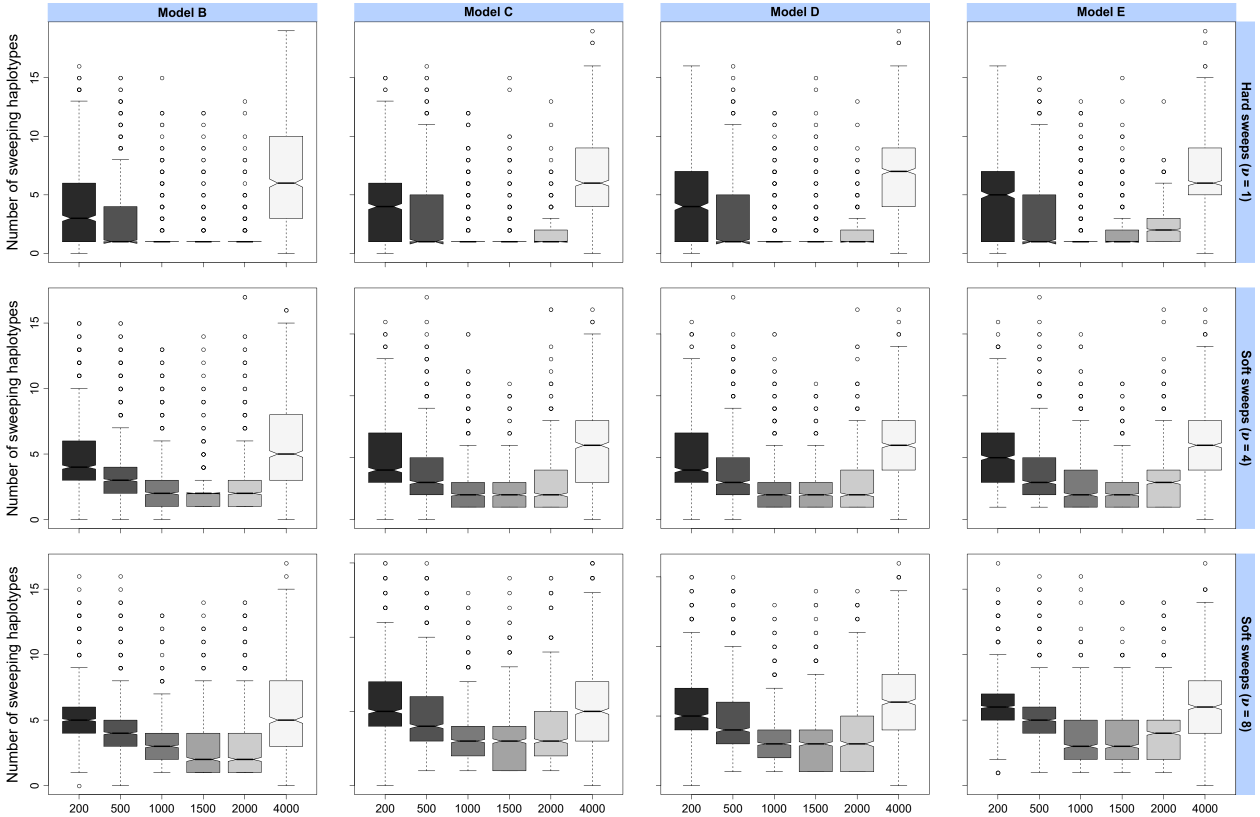


Figure S13: Box plots summarizing the distributions of the inferred number of sweeping haplotypes $\widehat{m}$ for different distortion variants (choosing alternate $f_i$; see *Theory*) under the European CEU human demographic model. Variant names are identical to Figure S12, with $f_i = (1/i)/\sum_{j=1}^{m} 1/j$ (model B), $f_i = (1/i^2)/\sum_{j=1}^{m} 1/j^2$ (model C), $f_i = e^{-i}/\sum_{j=1}^{m} e^{-j}$ (model D), and $f_i = e^{-i^2}/\sum_{j=1}^{m} e^{-j^2}$ (model E). Simulated selective sweeps were of strength $s \in [0.005, 0.5]$ drawn uniformly at random on a log-scale and drawn from $\nu \in \{1, 4, 8\}$ distinct sweeping haplotypes. $K = 20$ haplotype frequency spectra were used for inference, and data were identical to Figure S12.

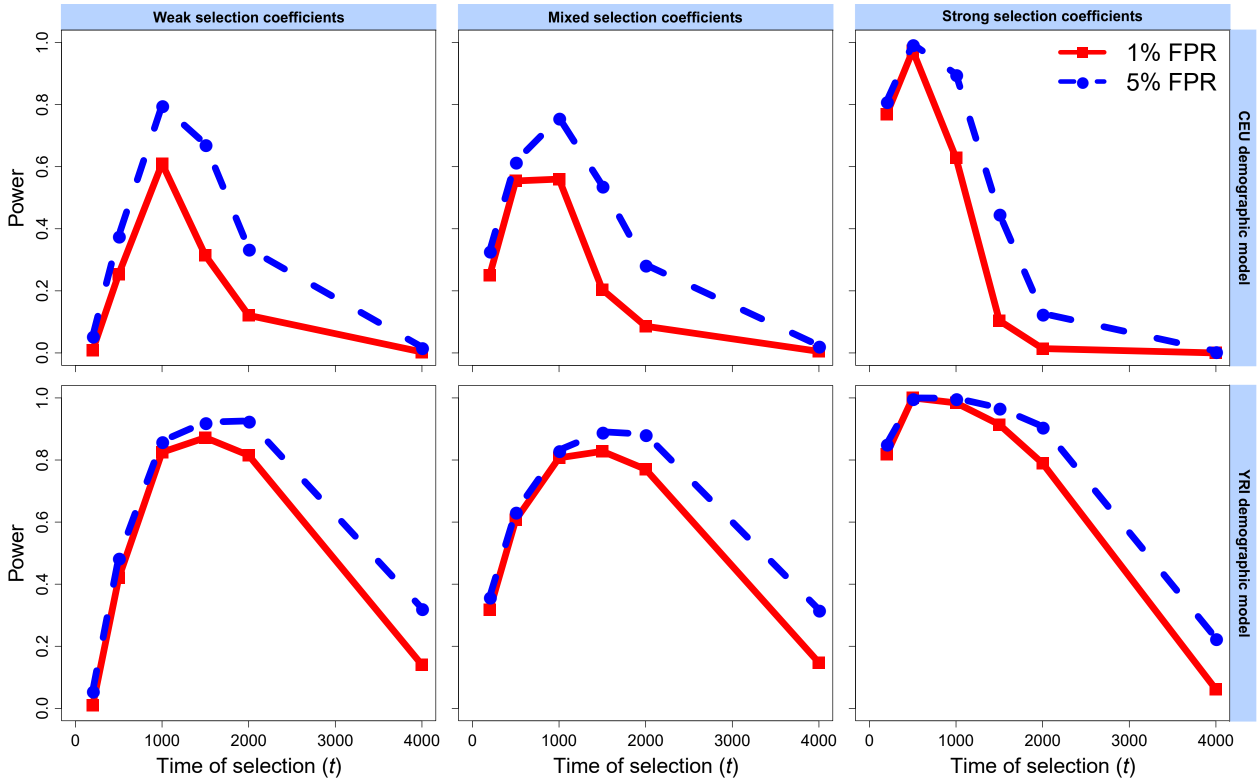


Figure S14: Power of H12 to distinguish simulated hard selective sweeps from neutrality at 1% and 5% false positive rates (FPRs). Selection begins at time $t \in \{200, 500, 1000, 1500, 2000, 4000\}$ generations before sampling under the European CEU (top) and sub-Saharan African YRI (bottom) human demographic models. Selection coefficients for sweep simulations were drawn uniformly at random from $s \in [0.005, 0.05]$ (weak coefficients, left), $s \in [0.005, 0.5]$ (mixed coefficients, middle), or $s \in [0.05, 0.5]$ (strong coefficients, right), and specifically drawn from a log scale for mixed sweeps. Simulated replicates are identical to those in Figure 2. All inferences used a spectrum of $K = 20$ for likelihood computations.

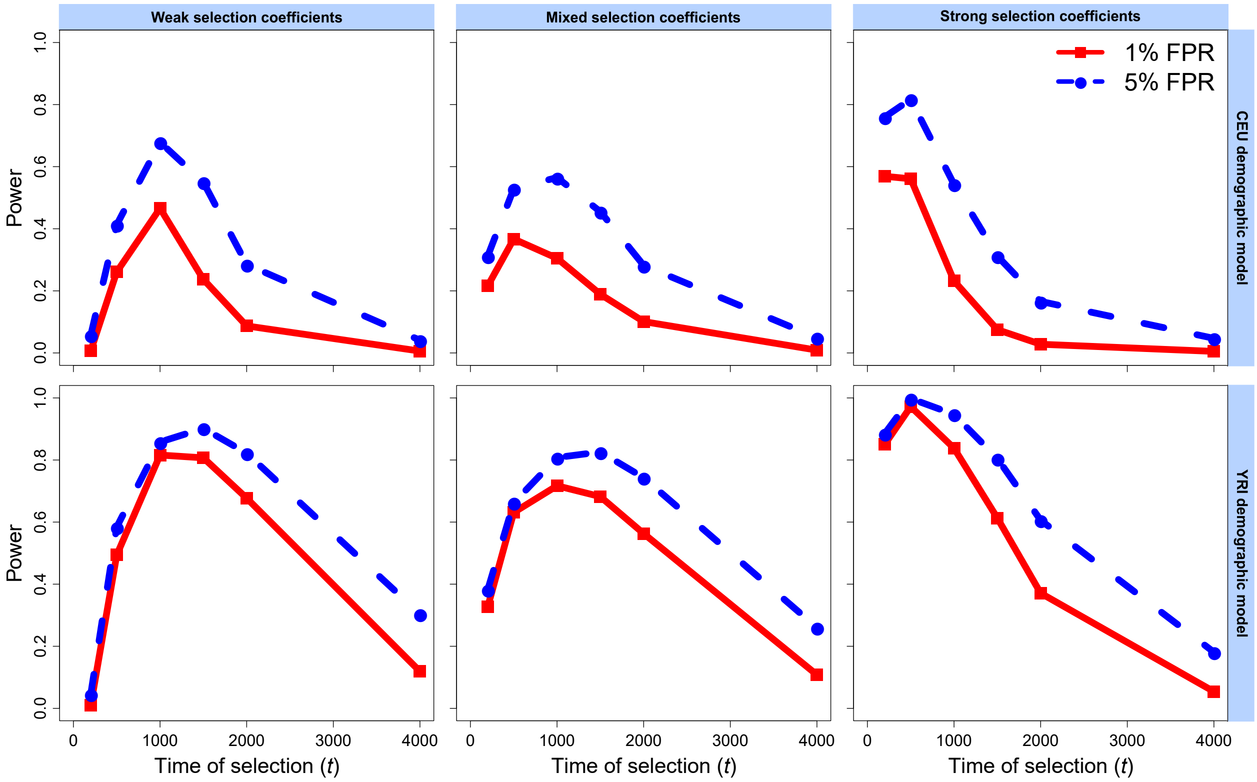


Figure S15: Power of H12 to distinguish simulated soft selective sweeps on four initially-selected haplotypes ($\nu = 4$) from neutrality at 1% and 5% false positive rates (FPRs). Selection begins at time $t \in \{200, 500, 1000, 1500, 2000, 4000\}$ generations before sampling under the European CEU (top) and sub-Saharan African YRI (bottom) human demographic models. Selection coefficients for sweep simulations were drawn uniformly at random from $s \in [0.005, 0.05]$ (weak coefficients, left), $s \in [0.005, 0.5]$ (mixed coefficients, middle), or $s \in [0.05, 0.5]$ (strong coefficients, right), and specifically drawn from a log scale for mixed sweeps. Simulated replicates are identical to those in Figure 3. All inferences used a spectrum of $K = 20$ for likelihood computations.

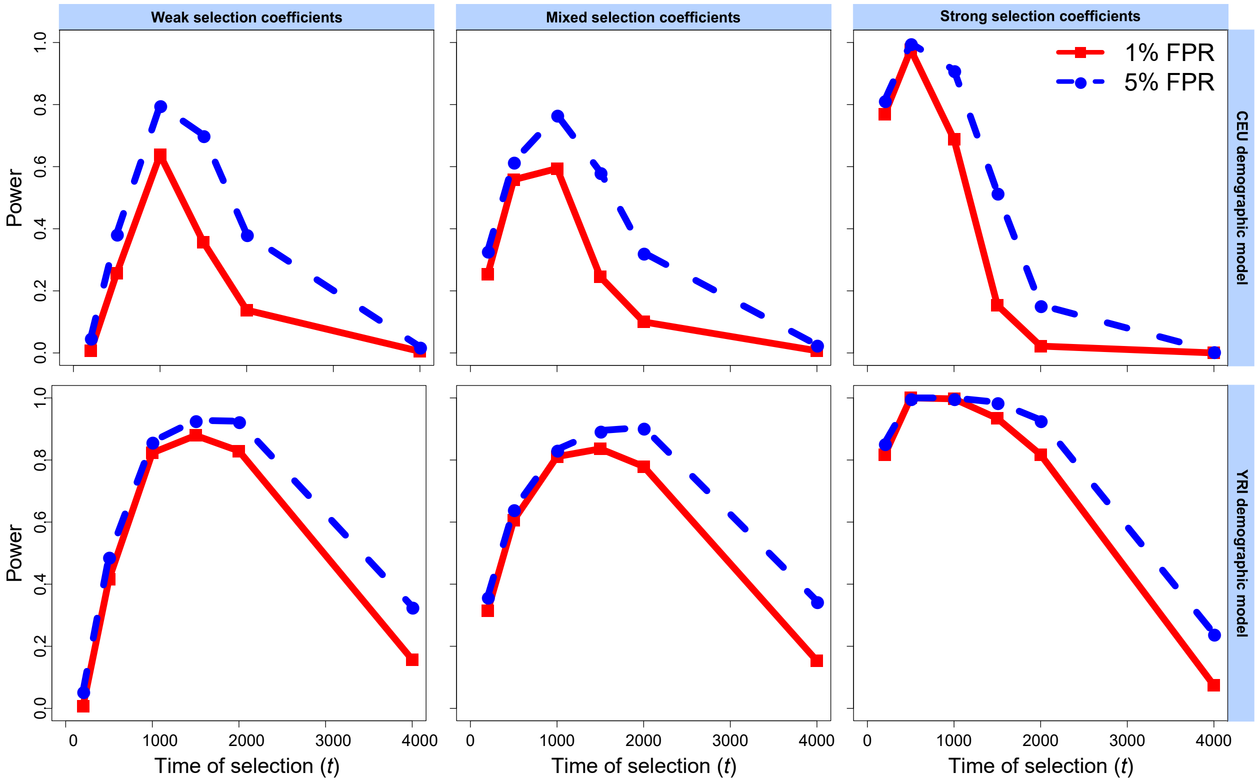


Figure S16: Power of G123 to distinguish simulated hard selective sweeps from neutrality at 1% and 5% false positive rates (FPRs). Selection begins at time $t \in \{200, 500, 1000, 1500, 2000, 4000\}$ generations before sampling under the European CEU (top) and sub-Saharan African YRI (bottom) human demographic models. Selection coefficients for sweep simulations were drawn uniformly at random from $s \in [0.005, 0.05]$ (weak coefficients, left), $s \in [0.005, 0.5]$ (mixed coefficients, middle), or $s \in [0.05, 0.5]$ (strong coefficients, right), and specifically drawn from a log scale for mixed sweeps. Simulated replicates are identical to those in Figure S7. All inferences used a spectrum of $K = 20$ for likelihood computations.

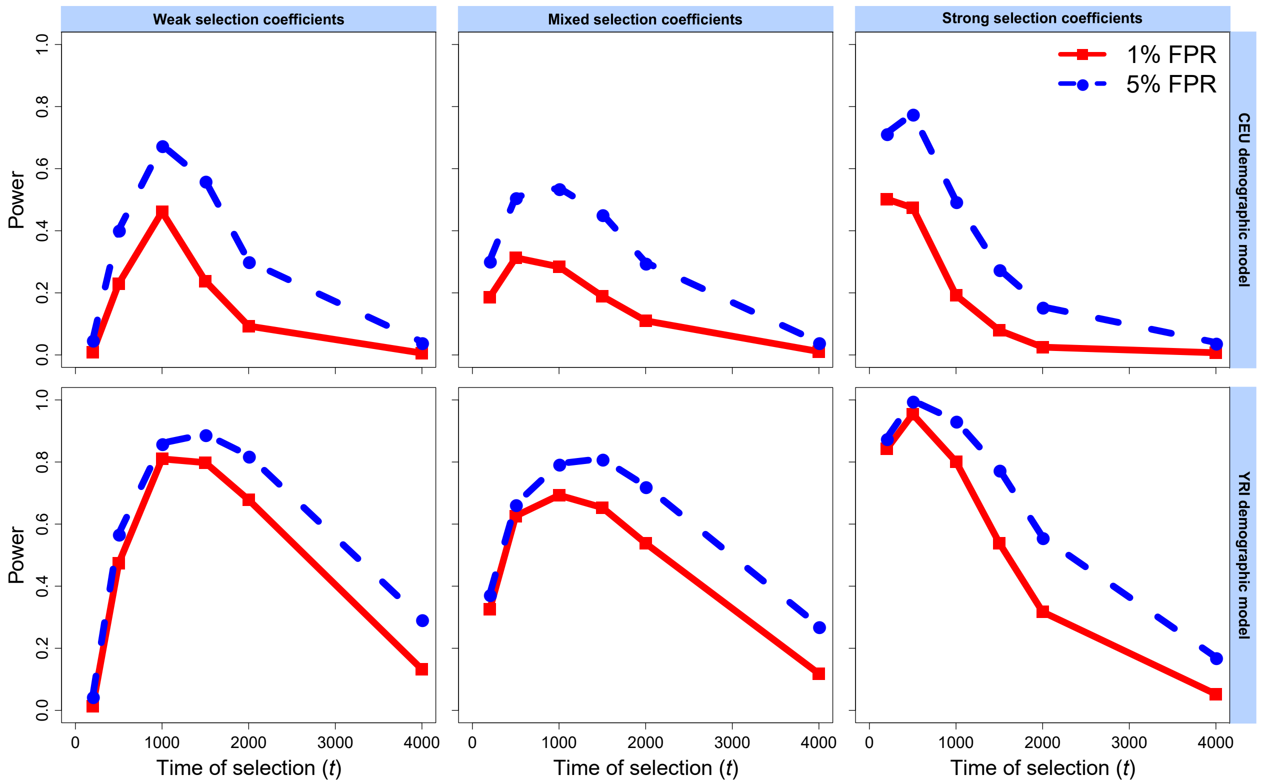


Figure S17: Power of G123 to distinguish simulated soft selective sweeps on four initially-selected haplotypes ($\nu = 4$) from neutrality at 1% and 5% false positive rates (FPRs). Selection begins at time $t \in \{200, 500, 1000, 1500, 2000, 4000\}$ generations before sampling under the European CEU (top) and sub-Saharan African YRI (bottom) human demographic models. Selection coefficients for sweep simulations were drawn uniformly at random from $s \in [0.005, 0.05]$ (weak coefficients, left), $s \in [0.005, 0.5]$ (mixed coefficients, middle), or $s \in [0.05, 0.5]$ (strong coefficients, right), and specifically drawn from a log scale for mixed sweeps. Simulated replicates are identical to those in Figure S8. All inferences used a spectrum of $K = 20$ for likelihood computations.
